# Behavioral and Neurostructural changes associated with Chronic Amygdala Hyperactivation

**DOI:** 10.1101/2021.09.11.459894

**Authors:** Keith A. Misquitta, Sierra A. Codeluppi, Jaime K. Knoch, Yashika Bansal, Toshi Tomoda, Jacob Ellegood, Jason P. Lerch, Etienne Sibille, Yuliya S. Nikolova, Mounira Banasr

## Abstract

**Background:** The amygdala (AMY) is a key brain region of the limbic system that plays a critical role in emotion processing and stress response. Functional magnetic resonance imaging (fMRI) studies identified abnormal AMY activation in psychiatric illnesses including major depressive disorder (MDD). Stress exposure is a major precipitating factor of MDD episodes which are associated with AMY hyperactivity. Preclinical studies using of pharmacologic, opto- and chemogenetic approaches to activate AMY neurons have consistently demonstrated that acute AMY hyperactivation induces anxiety-like behaviors in mice. However, it remains unknown if chronic hyperactivation of the amygdala (cHOA) is sufficient to induce chronic stress-like deficits or is a susceptibility factor for chronic stress-induced behavioral, volumetric and synaptic deficits.

**Methods:** Using designer receptor exclusively activated by designer drug (DREADD) approach, basolateral amygdala (BLA) neurons of Camk2a-cre mice infected with a virus driving the expression of the Gq-coupled DREADD were activated with clozapine-N-oxide (in drink water for 5 weeks). Mice were then exposed to chronic restraint stress (CRS; 1X/day for 1hr) for 2 weeks. All mice were behaviorally assessed in the Phenotyper (PT), and sucrose consumption tests (SCT) each week and in the novelty supressed feeding (NSF, once at the end of the experiment). Animals were then perfused for *ex vivo*-MRI and puncta density analysis.

**Results:** We found that mice with cHOA displayed a progressive increase in baseline anxiety-like deficits in the PT test and slightly more marked deficits following CRS compared to controls, but not statistically different from animals subjected to CRS alone. Also, cHOA did not exacerbate CRS effect in the NSF. No significant cAH effect was found in the SCT before or after CRS. MRI analysis revealed no statistical charges between groups, while increased synaptic puncta density was found in cHOA mice subjected to CRS compared to cHOA or CRS alone.

**Conclusion:** We demonstrate that cAH is sufficient to induce anxiety and may exacerbate CRS effects on anxiety and synaptic measures. Results also suggest that cHOA was not sufficient to induce depressive-like behavior and was not a vulnerability factor for stress-induced depressive-like behavior in mice. Altogether, our findings imply that a strong causal link between AMY hyperactivity and elevated anxiety, but not depressive-like behaviors and provide critical information to clinical research focused on using AMY activity level as a biomarker in stress-related illnesses.

## 1. Introduction

Major depressive disorder (**MDD**) is a major societal burden affecting over 264 million people worldwide, where patients suffer from prolonged and chronic emotional dysregulation (James et al., 2018). MDD is a multi-symptomatic illness often comorbid with anxiety symptoms and anxiety disorders (Groen et al., 2020; Keers and Aitchison, 2010; Najt et al., 2011). Although MDD and anxiety disorders are categorized separately in the DSM-V they do share overlapping clinical populations highlighting potential pathophysiological associations and shared underlying neurobiological mechanisms. Moreover, studies have shown that individuals with history of general anxiety disorders (**GAD**) recently (Patten et al., 2015) or in early childhood (Moffitt et al., 2007) are more likely to develop MDD, suggesting that chronic anxiety can lead to depression later in life.

Neuroimaging studies have identified brain-wide dysfunctions associated with MDD and anxiety disorders including within corticolimbic brain regions such as the amygdala (**AMY**) (McKinnon et al., 2009; Schmaal et al., 2017; Schmaal et al., 2020; Schmaal et al., 2016; Treadway et al., 2015). The AMY is an evolutionarily conserved brain structure across mammalians (Price, 2003), which plays a significant role in internal and external stimuli integration (LeDoux, 2003) and procession of emotionally salient information involved with memory (Phelps, 2004), decision-making (Gupta et al., 2011), emotional reaction (Ressler, 2010) and reward (Stuber et al., 2011). Magnetic resonance imaging (MRI) studies have identified inconsistent AMY volume changes in patients diagnosed MDD and anxiety disorders (Arnone et al., 2012a; Frodl et al., 2003; Hamilton et al., 2008; Schmaal et al., 2016; Van Tol et al., 2010). However, functional MRI (fMRI) studies consistently report hyperactivity of the AMY in similar clinical populations when patients are presented facial expressions with negative valence (Blair et al., 2008; Brühl et al., 2014a; Peluso et al., 2009). In MDD, amygdala hyperactivity was shown to be associated with symptom severity (Gaffrey et al., 2011; Goldin et al., 2009; Peluso et al., 2009; Wang et al., 2008) and symptom occurrence across age (Gaffrey et al., 2011; Tao et al., 2012; Yang et al., 2010a) in both sexes (Bremner et al., 2005; Yang et al., 2010a). Further amygdala hyperactivity is persistent following remission in unmedicated MDD patients (Victor et al., 2010) and is reduced following antidepressant treatment (Arnone et al., 2012b; Sheline et al., 2001; Victor et al., 2010). Following on this line of thought, studies have suggested that amygdala hyperactivity could be used as an biomarker to clinical and subclinical anxiety (Etkin et al., 2004; Etkin and Wager, 2007), as a predictor of vulnerability for MDD (Monk et al., 2008; Ramel et al., 2007) and future life stressors (Swartz et al., 2015), or even as an indicator of response to antidepressant treatment (Williams et al., 2015).

Stress exposure is a major influential factor to the onset of various neuropsychiatric illnesses including MDD and anxiety disorders that disrupts the homeostatic balance (Kessler, 1997; Kessler et al., 2003; McEwen, 2017). Stress-based animal models have been instrumental to understand the pathophysiology of MDD and anxiety. Chronic stress exposure induces behavioral and cellular deficits in rodents that exhibit maladaptive responses relevant to investigating MDD pathophysiology (Banasr et al., 2021; Nollet et al., 2013; Planchez et al., 2019). Recent chronic stress MRI studies in mouse and rat have identified volumetric changes to corticolimbic brain regions including the prefrontal cortex, hippocampus and AMY (Anacker et al., 2016; Kassem et al., 2013; Lee et al., 2009; Magalhães et al., 2018; Misquitta et al., 2021; Nikolova et al., 2018). Interestingly we recently demonstrated that while AMY volume changes following chronic stress seems inconsistent across models (unpredictable chronic mild stress (UCMS) and chronic restraint stress (CRS), structural covariance changes associated with chronic stress in the AMY can be found in both models (Misquitta et al., 2021; Nikolova et al., 2018). Indeed, structural covariance network (SCN) analysis conducted in both these studies identified an increase in structural connectivity of the AMY following chronic stress. Further this AMY SCN pattern reorganisation was also identified in young adults with a history of childhood trauma showing both MDD and anxiety symptoms (Nikolova et al., 2018).

Preclinical and clinical studies attribute neuroanatomical structural changes to cytoachitectural alterations including spine number changes or dendritic reorganisation. Chronic stress induces increased spine number (Hill et al., 2011), dendritic arborization and branching complexity of basolateral amygdala (BLA) neurons (Vyas et al., 2004). Recently, we linked chronic stress-induced AMY volume and strengthening of AMY SCN to puncta density of synaptic markers in the BLA (Misquitta et al., 2021; Nikolova et al., 2018), which also correlated with elevated emotionality behavior, notably anxiety. These finding highlight the role of BLA synaptic changes in the structural and behavioral deficits associated with chronic stress.

The BLA predominantly contains glutamatergic excitatory neurons (80-90%) and a relatively small population of GABAergic interneurons and neuroglia cells (Duvarci and Pare, 2014; Sharp, 2017; Spampanato et al., 2011). Acute activation of the BLA glutamergic neurons was shown to induce in anxiogenic response as observed in pharmacologic or chemo/optogenetic studies (Biselli et al., 2021; Hare and Duman, 2020; Muir et al., 2019; Siuda et al., 2016). Selective manipulation of excitatory projections specific to and from the BLA can results in either anxiogenic or anxiolytic responses acutely (Janak and Tye, 2015). Altogether, these studies focused on the understanding of the behavioral or cellular consequences of acute BLA excitatory neuron activation while the effects of chronic activation remain unknown. This question is of a particular interest since chronic stress is associated with chronic or sustained amygdala activation (Fee et al., 2020; Perrotti, 2004).

In this study we first determined if chronic hyperactivation of the AMY (**cHOA**) using a chemogenetic designer receptor exclusively activated by designer drugs (DREADD) approach to activate BLA glutamatergic neurons will lead to depressive-like behavior. To do so, we measured weekly cHOA changes in physical state, anxiety- and anhedonia-like behaviors for 5 weeks. We also assessed the behavioral effects of prior cHOA on the response to subchronic (3 days) or chronic stress (2 weeks). We then examined if cHOA alone or combined with chronic stress resulted in changes to AMY volume using *ex-vivo* MRI, and extended our structural analysis to the AMY circuit, regions involved MDD and generally using unbiased whole brain volumetric analysis. Finally, we examine potential association between cHOA, behavior or AMY volume with local synaptic changes quantified following immunohistochemistry.

## 2. Methods

### 2.1 Animals

Transgenic animals were generated from adult male homozygote CAMK2a cre/cre (Jackson Laboratory, Bar Harbor, Maine, USA, B6.Cg-Tg(Camk2a-cre)T29-1Stl/J, 005359) mice bred in house with female C57BL/6 mice (Jackson Laboratory, Bar harbor, Maine, USA). A total of 48 mice (50% females) F1 generation heterozygote CAMK2a cre/+ mice, 8-12 weeks old, were used in this study. Mice were housed under a 12-hour light/dark cycle and *ad libitum* food and water, except special water was provided or when specific behavioral testing required deprivation. The procedures were preformed according to Center for Addiction and Mental Health animal facility and Canadian Council on Animal Care (CCAC) guidelines.

### 2.2 Viral infusion, drug administration and chronic restraint stress

Mice anaesthetized using isoflurane were bilaterally infused with pAAV2-hSyn-DIO-hM3D(Gq)- mCherry (≥ 6 × 10^12^ vg/mL; Addgene #44361, Lot v58216). After exposing the skull and creating small <2mm bilateral holes with a dental drill (Stoelting, Cat# 58610), the viral infusion was performed using glass capillaries (outside diameter 20-50um) prepared using Gravipull-3 Micropipette Puller (Kation Scientific, LLC., Minneapolis, MN, USA) and lowered into the BLA (AP −1.3, ±3.0, DV −5.2 from bregma). The virus (0.3 ul/side) was delivered at a continuous rate of 0.1µl/min using a GenieTouch Syringe PumpPro (Kent Scientific Co., Torrington, CT, USA), followed by 5 min of wait time to allow diffusion. Analgesic (Rimadyl®, Fort Dodge Animal Health, Overland Park, KS; 5mg/kg) and topical antiseptic were administrated for 3 days as post-operative care. All procedures were conducted in aseptic conditions. Behavioral testing and drug administration began 4 weeks after surgery.

Clozapine-N-oxide (CNO, Hello Bio, United States, Cat#HB6149) was provided to half of the animals (n=24; 1:1 ratio of males and females) for 7 weeks at a dose of ∼5 mg/kg diluted in drinking water (Vetere et al., 2017), based on ∼8 mL/day fluid consumption in opaque bottles. Mice received freshly made CNO every 2-3 days to prevent deterioration due to light exposure or room temperature. The other half of the animals (n=24) i.e. mouse groups not treated with CNO were given normal drinking water (vehicle) in similar bottles at the same time.

Following 5 weeks of chronic CNO or vehicle administration, animal groups were split into 2 subgroups each, exposed or not to CRS (n=12/group; 1:1 male/female). CRS consist in placing each mouse inside a 50mL Falcon^®^ tube perforated with holes on either side for air flow, for a 1-hour period, once a day for 15 days.

### 2.3 PhenoTyper *Test*

For this test, each mouse was placed in a home cage-like apparatus or PhenoTyper (Noldus, Leesburg VA, USA) which is able to video-track using EthoVision (EthoVision 10 software) hourly time spend in two designated zones: one delineated in front of the food hopper (or food zone [6.5×15 cm]) and one delineated containing a covered shelter (shelter zone [10×10cm]). Time spent in each zone is recorded from 7:00PM to 07:00AM during the animal’s dark cycle. Between 11:00-12:00AM, an anxiogenic challenge i.e. spotlight over the food zone is applied for 1hour. Testing in the PhenoTyper box was performed weekly on days 0, 7, 14, 21, 28, 35 of the CNO treatment and on day 3 and day 15 of the CRS exposure. In previous studies, we identified that CRS-exposed and control animals spend equivalent amounts of time avoiding the food zone in favor of the shelter during the spotlight challenge. However, CRS animals subjected to chronic stress continue to avoid the food zone after the challenge (Codeluppi et al., 2021; Misquitta et al., 2021; Prevot et al., 2019). This is a behavior, is highly specific of chronic stress exposure, defined as “residual avoidance” (RA). Shelter and food zone RA were calculated from the difference between the time spent during the light challenge and the sum of the time spent avoiding the lit zone (5 hours post-challenge), normalized to the control (vehicle no stress group). Positive RA value indicates the animal’s avoidance of the lit zone in favor of the shelter zone (Prevot et al., 2019). Formulae for RA for each mouse as followed:

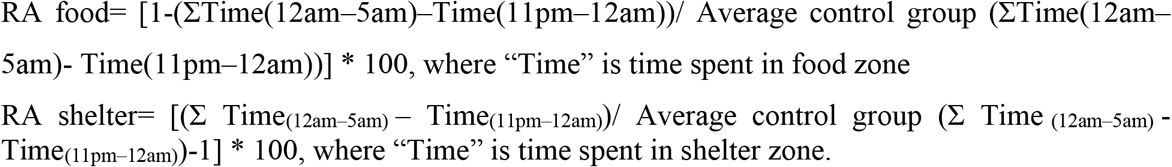

### 2.4 Sucrose Consumption Test (SCT)

For the first SPT, animals were habituated to a 1% sucrose solution (Sigma, MO, USA) for 48 hrs and then fluid deprived for 16 hours overnight. The next day, bottles were placed back on the cages at 10am, and sucrose consumption was measured for a 1-hour period. Bottles with no sucrose are then given to the mice and the same procedure (fluid deprivation and 1h test) is performed the next day with water to control for fluid intake (Codeluppi et al., 2021; Duric et al., 2017; Misquitta et al., 2021). Sucrose preference is calculated as percent of the ratio of sucrose consumption over total fluid consumption of the two 1hr-test. The entire procedure was performed every following week for 5 weeks, except that mice were only provided 24 hour period of habituation to the 1% sucrose solution. Sucrose consumption test occurred every week the day after the phenotyper test. However, to allow completing of testing (control fluid intake, phenotyper test, novelty suppressed feeding and locomotor activity) before perfusion of the animals at day 15, the last sucrose preference occurred on day 10 of the CRS exposure. CNO was provided in the sucrose and water bottles throughout the experiment.

### 2.5 Coat State and Weight Measurements

Every week before the phenotyper test, an experimenter blinded to the group assignment evaluated the coat state of each mouse by assessing, with a value of either 0 (well maintained), 0.5 (intermediary state) or 1 (unkept), seven anatomical regions of the mouse (head, neck, dorsal/ventral coat, forepaws, hindpaw and tail). The sum of values for each mice correspond to the coat state score (Nollet et al., 2013). Along with coat state assessment, animal weight (in grams) was recorded to assess weight gain or loss due to the manipulations.

### 2.6 Novelty Suppressed Feeding (NSF)

Prior to testing, mice were food deprived for 16 hours. Mice are then placed in dimly lit (lux 28-30) novel arena (45 × 30 × 27 cm) with access to a single food pellet position in the center of the arena. Latency to bite (in sec) the pellet was measured by an experimenter blinded to the group assignment. To control for potential biases associated with appetite drive, a similar test was performed in the animal’s home cage. NSF occurred once at the end of the experiment on day 13 of CRS.

### 2.7 Locomotor activity

On day 14 of CRS, mice were individually placed in new cages identical to their home cage and recorded using a camera on the ceiling positioned over the cages. Using Anymaze (Stoelting Co., Wooddale, IL,USA) tracking software, total distance travelled (in meters) is measured for a 30 min.

### 2.8 Mouse Brain Tissue Preparation and MRI

Tissue preparation and MRI will be performed as previously described (Misquitta et al., 2021; Nikolova et al., 2018). Briefly, mice anesthetized with avertine (125mg/kg, i.p.) will be intracardially perfused at a rate of approximately 100mL/hr with 30 ml of 0.1M phosphate-buffered saline (PBS) containing heparin (10U/mL) and ProHance (2mM, Bracco Diagnostics, NJ, USA), followed by 30 ml of 4% paraformaldehyde solution containing 2mM ProHance. After removal of surrounding tissues (skin, muscles, lower jaw, etc), brains left intact within the skull are post-fixed in PFA (4% + 2mM ProHance) overnight at 4°C. Samples were then transferred into a buffer storage solution (0.02% sodium azide + 2mM ProHance) for a minimum of one month (De Guzman et al., 2016) before MRI using a 7.0 Tesla MRI scanner (Agilent Inc., Palo Alto, CA). Scanning was performed as in Misquitta et al. (2021) using a regional volume parcellation into 182 brain regions identical to a previous study (Misquitta et al 2021). We analysed region of interest (ROI) volumes using averages across hemispheres to minimize multiple testing burden. Absolute brain volumes (in mm^3^) were determined using deformation-based morphometry. First, we conducted 2 initial *a priori* analyses focusing on the AMY volume/AMY circuit or on 26 brain regions pre-selected for involvement in MDD (Misquitta et al., 2021). The analysis of the AMY-related circuit required summation of several subregions for the hippocampal formation (CA1, CA2, CA3 and dentate gyrus), the anterior cingulate cortex (area 24 a,a’,b,b’), prelimbic (cingulate cortex area 30/32), infralimbic (cingulate cortex area 25) and cingulate cortex area 29 (area a, b, c). Finally, we conducted a whole brain analysis employing all 182 ROIs i.e. without these summations.

### 2.9 Fluorescence immunohistochemistry and puncta analysis

Following ex vivo MRI, brains removed from skulls were cryoprotected in PBS-30% sucrose. Thirty μm thick free-floating sections were cut using a cryostat (Leica, Wezlar, German), stored in at −20°C in sucrose 30%, polyvinyl-pyrrolidone-40 (1%), 0.1M phosphate buffer (PB) and ethylene glycol 30%) for cryoprotection. Sections containing the BLA (coordinates from Bregma − 1.31 and − 1.43 mm) were used for immunohistochemistry. After several washes in 1X PBS at 4°C and incubation in an antigen retrieval solution (0.01M sodium citrate) at 80 °C for 15 mins, sections were cooled to room temperature. Sections were then placed in 0.3% X-100-Triton (20 min), followed by incubation in 20% donkey serum (diluted in PBS) 1hr. Next, sections were placed overnight at 4°C in a solution containing 2% donkey serum and primary antibodies for post-synaptic density protein 95 (PSD95; rabbit host, 1:100, Cell Signaling Technology, Danvers, MA, product #2507, lot 2), vesicular glutamate transporter 1 (VGLUT1; guinea pig host, 1:500, Synaptic Systems, Goettingen, Germany, product #135304, lot 2-57) and mCherry (chicken host, 1:1200, Abcam, Cambridge, UK, product # ab 205402, lot# GR3258636-1). After a PBS wash, sections were incubated for 2 h in 2% donkey serum solution containing secondary antibodies conjugated to anti-rabbit Alexa488 (Invitrogen, Massachusetts, USA, CAT#A21206, Lot#1834802) and anti-guinea pig CF405M (Biotium, Hayward, CA, CAT# 30376, Lot# 14C023), and anti-chicken Alexa568 (Invitrogen, Massachusetts, USA, A11041, Lot#1963088) at 4°C to detect PSD95, VGLUT1 and mCherry immunoreactivity, respectively (1:500 for all). After a final wash in PBS (20min), sections were mounted onto Superfrost plus gold slides (Fisher Scientific, Massachusetts, United States), coversliped (Vectashield Antifade Mounting Media, Vector Laboratories, Burlingame, CA) and stored at 4°C until imaging. Image acquisition and deconvolution was performed as previously described (Misquitta et al., 2021). PSD95 and VGLUT1 puncta analysis was conducted using IMARIS v9.5.1 software (BitPlane, Badenerstrasse, Zurich, Switzerland) with quality parameters, i.e. threshold for the intensity of light at the center of each spot set at a quality score 300 for PSD95 and 350 for VGLUT1 (quality score). All results were calculated as the density in μm3.

Virus localisation in the BLA was verified using mCherry staining. five animals had to be excluded from the entire study including behavior, MRI and synaptic analysis for lack of adequate targeting. The n of each group is: vehicle (n=12; 6 males, 6 females), CNO (n= 9; 5 males, 4 females), vehicle+CRS (n= 12; 6 males, 6 females), CNO+CRS (n= 10; 4 males, 6 females).

### 2.10 Statistical analysis

Analysis of variance (ANOVA) was conducted on behavioral performances, AMY volume and synaptic data using SPSS statistical software (IBM, NY) with group, stress and sex as variable. Potential changes in behavioral performance assessed weekly i.e. the PhenoTyper test, sucrose consumption test, weight and coat state quality were assessed using repeated measures ANOVA with group, stress, sex and time as variable. To summarize performances across the multiple behavioral endpoint tests into one value, a z-emotionality score was calculated (Guilloux et al., 2011). The calculation normalizes the z-score of each test with respect to the mean and standard deviation of the control group of the respective sex. Z-emotionality score is an average of the individual test z-score. Z-emotionality calculation used the following measures: coat state, baseline and RA in PhenoTyper test and sucrose preference measured only in the last week of testing as well as the latency to bite in the NSF. For all aforementioned analysis, statistical significance was set at p ≤ 0.05. MRI analysis was conducted using ANCOVA using total brain volume (TBV) and sex as regressors and group, stress as factors. For the analysis of 26 MDD-associated and 182 brain regions, resulting p-values were corrected for multiple comparisons using false discovery rate (FDR, q<0.05). Pearson correlational analysis was performed to identify relationships between z-emotionality, and volume changes of AMY and AMY-associated regions as wells as the 26 MDD-associated regions. Lastly a Pearson correlation was performed to identify the relationship between synaptic puncta (PSD95 and VGLUT1) and z-emotionality or AMY volume.

## 3. Results

### 3.1 Longitudinal effects of cHOA in the phenotyper test

We used the PhenoTyper test to assess potential longitudinal of anxiety-like behaviors in animals with cHOA with or without CRS. Note that data from one vehicle male animal was not captured at the week 1 time point due to camera malfunction. Since data for this animal were collected every following week, we imputed the mean of the group for the time spent in the food and shelter zones at this time point to not remove the animal from the study.

Analysis of time in the food zone using repeated measures ANCOVA showed a significant main effect of drug (**Figure 1A-F**) on week 2 (F_(2, 492)_ = 4.09; p < 0.05), and week 5 (F_(2, 492)_ = 4.629; p < 0.05), with a trend towards significance on week 3 (F_(2, 492)_ = 3.168; p =0.08). Further a significant effect of drug x time interaction on week 1 (F_(12, 492)_ = 2.903; p < 0.001), week 2 (F_(12, 492)_ = 4.293; p < 0.0001), week 3 (F_(12, 492)_ = 2.343; p < 0.01) and week 5 (F_(12, 492)_ = 2.761; p < 0.01) with a trend on week 4 (F_(12, 492)_ = 1.693; p =0.065). Specifically, we observed a decrease in time spent in the food zone between 7:00pm to 11:00pm in mice with treated with CNO compared to vehicle group. This effect was significant on multiple hourly bins every week and observed predominantly during the first 4 hours of the test (baseline 7-11pm) before the light challenge (**Figure 1A-F)**.

**Figure 1:**
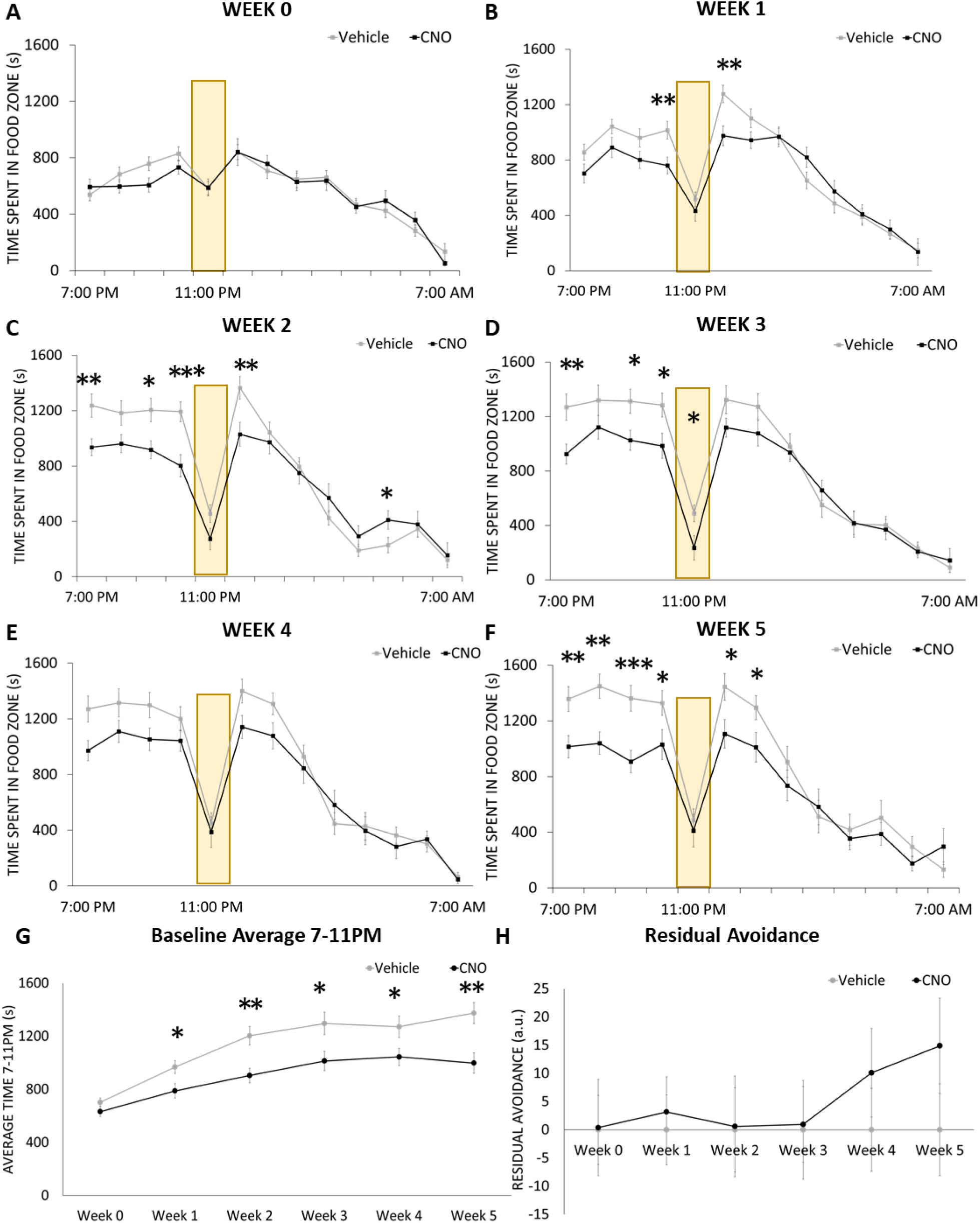
Longitudinal effects of chronic hyperactivation of the amygdala on anxiety-like behavior in the phenotyper test (food zone). Food zone time of mice treated or not with clozapine-N-oxide (CNO) was measured hourly between 7PM to 7AM in the PhenoTyper test. A white spotlight challenge occurs between 11-12PM (yellow bars). The time spent in the food zone prior to treatment (week 0, **A**) and every week of treatment (week 1, **B**; week 2 **C**, week 3, **D**; week 4, **E**; week 5, **F**) was assessed. Progressive changes in calculated baseline (7-11pm) averages and residual avoidance are shown in (**G)** and (**H)** respectively. Data collected for the shelter zone are in **Supplementary Figure 1**. Results are expressed as mean ± s.e.m. *p < 0.05, **p < 0.01, ***p<0.001 compared to vehicle treated group.

To further investigate this behavior, average of time spent in the food zone during the first 4 hours (baseline 7-11pm average) was calculated for each animal. Repeated measures ANOVA of baseline 7-11pm averages revealed a significant main effect of drug (F_(1, 205)_ = 11.161; p < 0.01; **Figure 1G**). This effect was explained by a significant decrease in time spent in the food zone in animals treated with CNO compared to vehicle on week 1 (p < 0.05) and every following week (week 2 (p < 0.01), week 3 (p < 0.05), week 4 (p < 0.05), week 5 (p < 0.0001). This demonstrated that animals with cHOA significantly spent less time in the food zone than controls. Sex contribution analysis revealed no significant main effect of sex, drug x sex, sex x time and drug x sex x time interaction on the weekly measures or on baseline 7-11PM measures (**Supplementary Figure 3**).

Repeated measures ANOVA of performances after light challenge measured by residual avoidance behavior revealing no significant main effect of drug or drug x time interaction (**Figure 1H**). Inclusion of sex as a factor showed no main effect of sex and no drug x sex interaction at any weekly time point (**Supplementary Figure 3**).

Analysis of time in the shelter zone using repeated measures ANOVA revealed no significant main effect of drug across the 5 time points. However, a significant main effect of drug was identified on week 2 (F_(1, 492)_ = 1.971; p < 0.05), weeks 3 (F_(1, 492)_ = 1.961; p < 0.05), 4 (F_(1, 492)_ = 2.12; p < 0.05) (**Supplementary Figure 1A-F)**, reflecting differences between CNO and vehicle groups at multiple hourly bins each week. Specifically, we found that mice with cHOA spend significant more time in the shelter than controls. Again, this effect was observed primarily during the first 4 hours of the test (baseline 7-11pm) before the light challenge. Inclusion of sex as a factor revealed a significant main effect of sex on week 4 (F_(1, 468)_ = 5.80; p < 0.05) and week 5 (F_(1, 468)_ = 12.057; p < 0.01) and drug x sex interaction on week 5 (F_(1, 468)_ = 4.135; p < 0.05). A sex x time interaction during week 0 (F_(12, 468)_ = 4.993; p < 0.0001), week 1 (F_(12, 468)_ = 1.829; p < 0.05), week 2 (F_(12, 468)_ = 2.877; p < 0.001), week 3 (F_(12, 468)_ = 2.873; p < 0.001), and week 4 (F_(12, 468)_ = 3.975; p < 0.0001) was also observed. Lastly, drug x sex x time interaction was identified on week 1 (F_(12, 468)_ = 1.816; p < 0.05) and week 5 (F_(12, 468)_ = 1.801; p < 0.05). The significant effects of sex are explained by female mice spending overall less time in the shelter following the light challenge. *Post-hoc* analysis showed no significant difference between individual groups for any hourly bins across the 5 time points.

Analysis of the average baseline 7-11PM behavior revealed a significant main effect of drug (F_(1, 205)_ = 12.77; p < 0.0001; **Supplementary Figure 1G**) and no significant drug x time interaction. Including sex as a fact showed a significant effect of drug on week 2 (F_(1, 46)_ = 9.519; p < 0.01), weeks 3 (F_(1, 46)_ = 5.451; p < 0.05), 4 (F_(1, 46)_ = 8.352; p < 0.01), 5 (F_(1, 47)_ = 10.444; p < 0.01), explained by female animals spending overall less time in the shelter but post hoc analysis showed that no difference between individual groups. No sex x time or drug x sex x time interactions were found (**Supplementary Figure 4)**.

Analysis of residual avoidance behavior after the light challenge over the 5 weeks of treatment revealed no significant main effect of drug, and drug x time interaction (**Supplementary Figure 1H)**, or sex (**Supplementary Figure 4)**.

### 3.2 Effects of prior cHOA and CRS response in the phenotyper test

Effects of 3 days or 15 days of CRS exposure in mice with or without prior cHOA was examined in the phenotyper test. At the 3 day-time point, repeated measures ANCOVA revealed a significant main effect of stress (**Figure 2A**; F_(1, 468)_ = 9.065; p < 0.01) but no significant main effect of drug, or drug x stress interaction. However, there was a significant stress x time interaction (F_(1, 468)_ = 2.424; p < 0.01) but no drug x time or drug stress x time interaction. This significant effect of stress is explained by CRS mice spending less time in the food zone after the light whether or not they were also treated with CNO, however post hoc analysis was unable to detect significant differences between group at any hourly bins. When splitting for sex there was no significant main effect sex and a stress x sex x time interaction.

**Figure 2:**
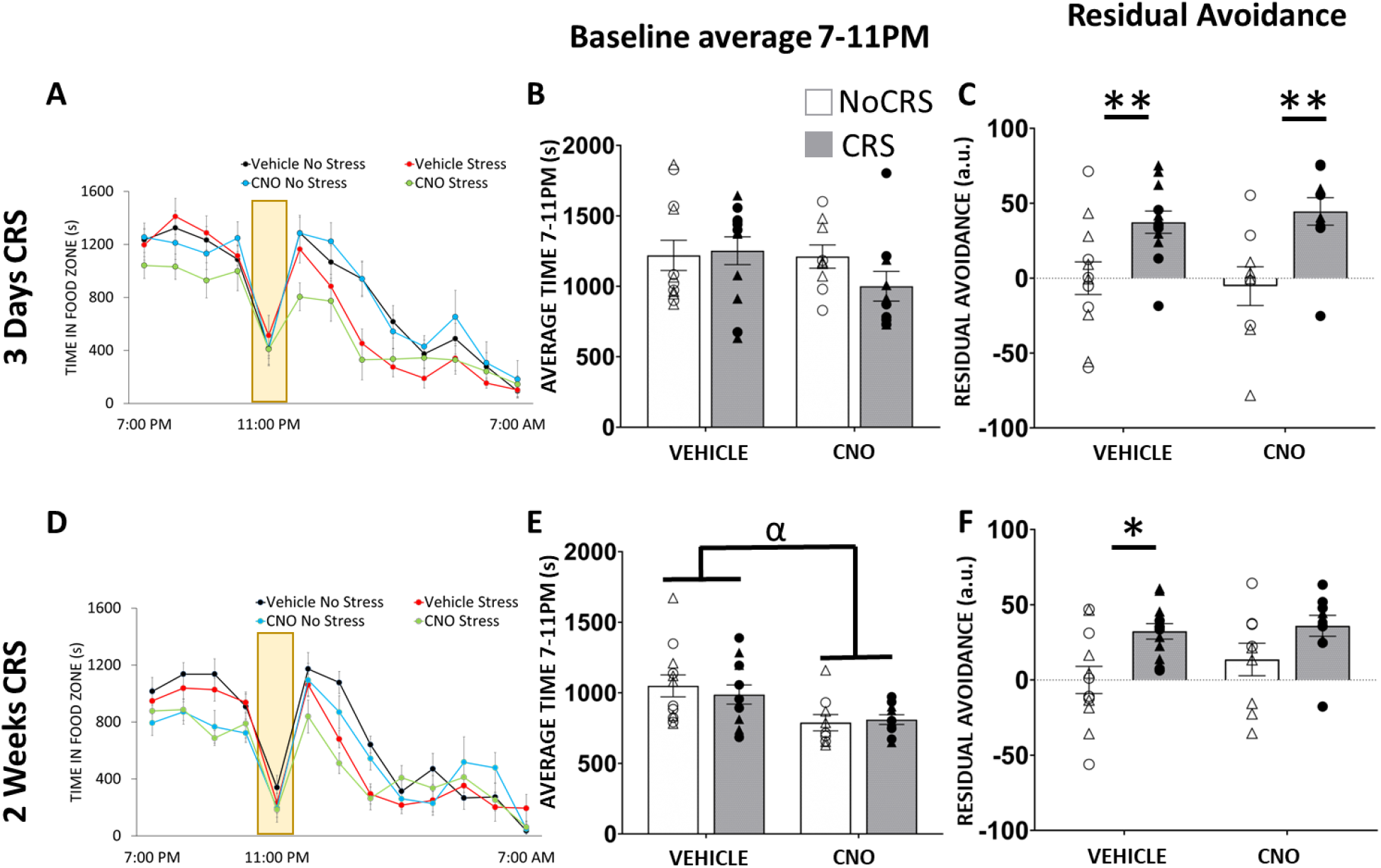
Effects of chronic hyperactivation of the amygdala combined or not with chronic stress on anxiety-like behavior in the phenotyper test (food zone). Food zone time of mice treated with vehicle, clozapine-N-oxide (CNO) and/or chronic restraint stress (CRS) was measured hourly between 7PM to 7AM in the PhenoTyper test on day 3 (**A**) and on day 15 of CRS exposure (**D**). A white spotlight challenge occurs between 11-12PM (yellow bars). Changes in calculated baseline (7-11pm) averages and residual avoidance found following 3 days of CRS are shown in (**B)** and (**C)** respectively. Baseline (7-11pm) averages (**E**) and residual avoidance (**F**) were also measured following 15 days of CRS. In B, C, E, F, Value collected for each mouse as well as sex is represented (Δ for males and O for females). Data collected for the shelter zone are in **Supplementary Figure 2**. Results are expressed as mean ± s.e.m. *p < 0.05, **p < 0.01 compared to vehicle treated group, α p<0.05, main effect of drug.

Average baseline (7-11pm) analysis revealed no significant main effect of drug, stress or drug x stress interaction (**Figure 2B)** or sex effects or sex x conditions interactions. Analysis of residual avoidance identified a significant stress (**Figure 2C;** F_(1, 39)_ = 18.479; p < 0.001), but no main effect of drug or drug x stress interaction, demonstrating that increased RA in CRS animals whether or not they were also treated with CNO. Inclusion of sex as a factor revealed no significant main effect of sex, drug x sex, stress x sex or drug x stress x sex interactions.

A similar analysis was conducted at the 15 day-time point **(Figure 2D-F)**, We found a significant main effect of drug (F_(1, 468)_ = 5.305; p < 0.05) and stress (F_(1, 468)_ = 4.96; p < 0.05) but no drug x stress interaction. There also was a drug x time (F_(1, 468)_ = 2.971; p < 0.001) and stress x time interactions (F_(1, 468)_ = 2.822; p < 0.01) but no drug x stress x time interaction. These effects were explained overall reduction of time spent in the food zone in cHOA mice during the 4 first hours of testing and reduction of time spent in food zone in CRS mice after the light challenge (**Figure 2D)**. However, post-hoc analysis was unable to detect significant differences between group at any hourly bins. Inclusion of sex as a factor revealed no main effect of sex but a significant stress x sex interaction (F_(1, 480)_ = 1.807; p < 0.05), explained by males showing greater reductions in time in the food zone compared to females after the challenge. Average baseline (7-11pm) analysis revealed a significant main effect of drug (**Figure 2E;** F_(1, 39)_ = 11.303; p < 0.01) but no significant main effect of stress or drug x stress interaction, explained by continued reduction of time spend in the food zone in cHOA animals. There was no significant effect of sex, drug x sex or stress x sex interactions on this measure. Analysis of residual avoidance revealed a significant main effect of stress (**Figure 2F;** F_(1, 39)_ = 11.405; p < 0.01) but no significant main effect of drug or drug x stress interaction, reflecting that CRS mice with or without cHOA spent less time in the food zone. There was no significant main effect of sex, no drug x sex and a significant stress x sex F_(1, 35)_ =4.416; p < 0.05) interaction explained by males showing greater reductions in time in the food zone compared to females after the challenge.

Similar analysis was performed for shelter time. At the 3 day-time point, we found a significant main effect of stress (**Supplementary Figure 2A;** F_(1, 468)_ = 15.232; p < 0.001)) but no significant main effect of drug, drug x stress, drug x time or drug x stress x time interaction. There was a significant stress x time interaction (F_(12, 468)_ = 4.6; p < 0.0001), reflecting that CRS mice with or without cHOA spent overall more time in the shelter zone. However, groups were not different in any specific hourly bins. Analysis of baseline 7-11PM averages of time spent in the shelter revealed no significant main effect of drug, stress and drug x stress interaction (**Supplementary Figure 2)**. Further, there was no main effect of sex, drug x sex, stress x sex and drug x stress x sex interaction. Analysis of residual avoidance in the shelter identified no significant main effect of stress, drug or drug x stress interaction (**Supplementary Figure 2)** or sex and sex x condition interactions.

At the 15 day-time point we found a significant stress x time interaction (**Supplementary Figure 2**; F_(12, 468)_ = 3.278; p < 0.001) but no significant main effect of drug, stress, drug x stress, drug x time and drug x stress x time interaction. Analysis of baseline 7-11PM averages of time spent in the shelter revealed no significant main effect of drug, stress or sex and no interactions between conditions (**Supplementary Figure 2**). Analysis of residual avoidance found a significant main effect of stress (**Supplementary Figure 2**; F_(1, 39)_ = 6.648; p < 0.05) but no significant drug or drug x stress interaction, reflecting that CRS mice with or without cHOA spent overall more time in the shelter zone, and no sex stress or interactions.

### 3.3 Effects of cHOA on the sucrose preference, coat state and weight

Coat state quality, sucrose preference and weight were assessed weekly to determine the effects of HOA alone. Repeated measures ANCOVA of sucrose preference revealed no significant main effect of drug or drug x time interaction (**Supplementary Figure 5**) or main effects of sex or interactions between sex and other factors (**Supplementary Figure 5**). Analysis of coat state scores revealed no significant drug or drug x time interaction for coat quality **(Supplementary Figure 5**). Including sex as a factor revealed a main effect of sex (F_(1, 195)_ = 22.426; p < 0.0001), a sex x time interaction (F_(5, 195)_ = 7.166; p < 0.0001) and no drug x sex x time interaction, reflecting that overall greater coat state degradation in male compared with female mice **(Supplementary Figure 5)**. Lastly, repeated measures ANCOVA of weight measures revealed no significant main effect of drug or drug x time **(Supplementary Figure 5**). Inclusion of sex in the analysis showed a main effect of sex (F_(1, 195)_ = 242.38; p < 0.0001) and a drug x sex x time interaction (F_(5, 195)_ = 2.958; p < 0.05), explained by overall greater weight in males compared with females **(Supplementary Figure 5)**.

### 3.4 Combined effects of cHOA and CRS in endpoint behaviors

Coat state assessment, sucrose preference and NSF tests were conducted once more in these animals following 2 weeks of CRS exposure not, to determine if cHOA effects are exacerbated by CRS. In the NSF test, we identified a significant main effect of stress (F_(1, 39)_ = 36.059; p < 0.0001; **Figure 3A**) but no significant effect of drug or drug x stress interaction. This effect was explained by a significantly increase in latency to bite in animals exposed to CRS with or without cHOA. No effects of sex or interactions were found on this measure. To control for appetite drive the latency to bite in home cage was measured. No difference between groups in home cage latency feed was found (data not shown).

**Figure 3:**
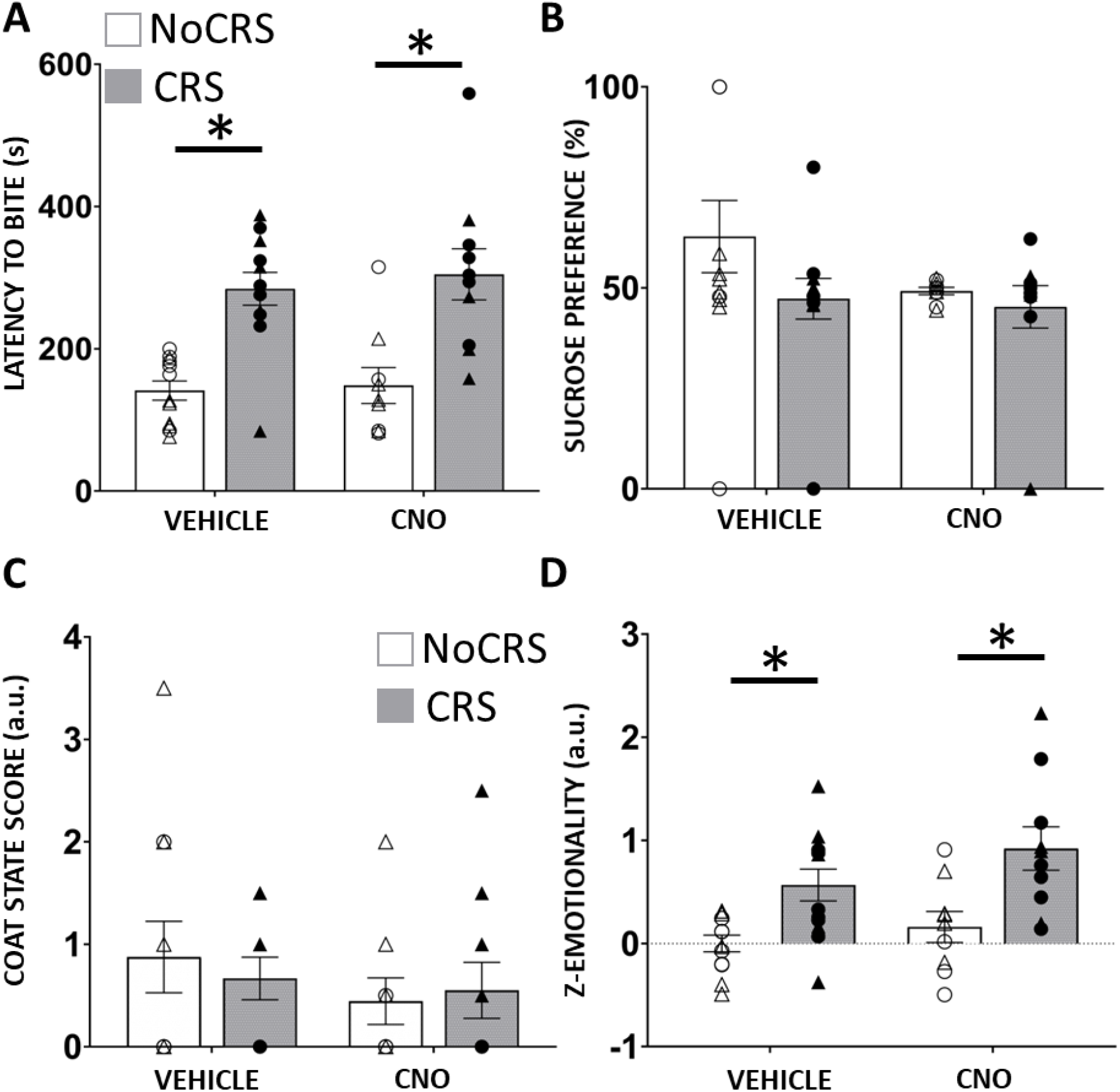
Effects of chronic hyperactivation of the amygdala combined or not with chronic stress on anxiety-, anhedonia-like behaviors, coat state and overall z-emotionality. Data from mice treated with vehicle, clozapine-N-oxide (CNO) and/or chronic restraint stress (CRS) were collected in the novelty suppressed feeding test (**A**), the sucrose preference test (**B**) and coat state assessment (**C**). Overall z-emotionality scores calculated with respect to average control group are shown in (**D**). Value collected for each mouse as well as sex is represented (Δ for males and O for females). Results are expressed as mean ± s.e.m. *p < 0.05 compared to vehicle-no CRS treated group.

Analysis of percent sucrose consumption revealed no significant main effect of drug, stress or drug x stress interaction **(Figure 3B)**. This remained unchanged when sex was added as a factor in the analysis. Analysis of coat state score revealed no significant main effect of drug, stress or drug X stress interaction **(Figure 3C)**. However, we found a only significant main effect of sex (F_(1, 35)_ = 12.436; p < 0.0001) where males show overall slightly greater coat state quality than females; however post hoc analysis found no significant difference between groups.

Performances in all tests conducted the last week of CRS i.e. NSF, percent sucrose consumption, coat state quality and phenotyper (baseline 7-11pm) were z-score normalized to control group average into a z-emotionality score. Analysis of z-emotionality revealed a significant main effect of drug (F_(1, 39)_ = 3.837; p < 0.05) and stress (F_(1, 39)_ = 33.653; p < 0.0001) but no drug x stress interaction **(Figure 3D)**. Post hoc analysis showed that both CRS groups i.e. with or without cHOA displayed significantly greater z-emotionality than no CRS groups. No significant effects of sex or interactions were found in this measure.

Finally, locomotor activity of all animals was measured and statistically analysis found no main effects of drug, stress, sex or conditions interactions (supplemental Figure.

### 3.5 Effects of cHOA and CRS on MRI-measured volume and synaptic changes

Structural adaptations induced by cHOA and CRS alone or combined were assessed using MRI analysis. We found no differences between groups in total brain volume (TBV) i.e no drug, stress, sex or interactions between factors. Further, Pearson correlation analysis identified no significant correlation between TBV and z-emotionality behavior. In an initial a priori analysis, we analyzed volumes of the AMY, BNST, PFC, ACC, infra-limbic, retrosplenial area, prelimbic area, hippocampal formation and nucleus accumbens (**Table**). Analysis conducted using TBV as a covariate and sex as a cofactor found no difference in volume between groups across these ROIs. Similarly, no significant volume changes between groups were identified cross the 26-MDD associated ROIs (**Supplementary Table 1**). Unbiased brain-wide analysis revealed no volumetric differences following cHOA, CRS alone or combined (**Supplementary Table 2**).

**Table 1:**
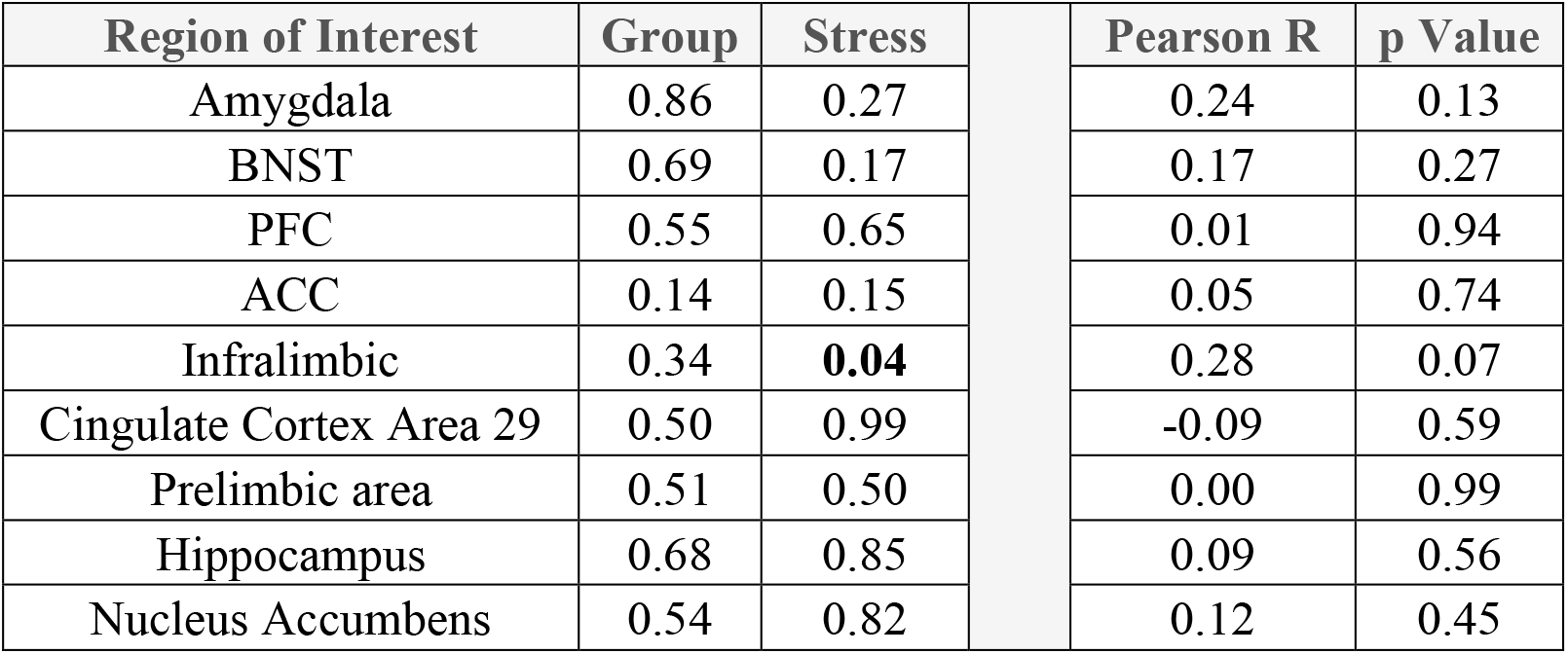
Analysis of variance of amygdala-associated brain region volumes and Pearson correlation with z-emotionality score.

We also analysed the cellular effects of cHOA and/or CRS at the synaptic level in the BLA using PSD95 and VGLUT1 puncta density. ANCOVA PSD95 puncta density identified no main effect of drug or sex but a significant drug x stress (F_(1, 39)_ = 4.218; p < 0.05, **Figure 4A**) Post hoc analysis revealed an increase in PSD95 puncta density between mice cHOA mice exposed CRS when compared to CRS and cHOA mice alone. Inclusion of sex as a factor revealed no main effects of sex or interactions with the other factors. Lastly, ANCOVA of VGLUT1 puncta density revealed a significant drug x stress (**Figure 4B**; F_(1, 39)_ = 4.49; p < 0.05) with no significant main effects of drug or stress. Post hoc analysis found an increase in VGLUT1 puncta density between mice cHOA mice exposed CRS when compared to CRS and cHOA mice alone. No significant effect of sex or interaction with other factors was identified. PSD95 and VGLUT1 puncta density did not correlate with z-emotionality and amygdala volume (Pearson r, p>0.05).

**Figure 4:**
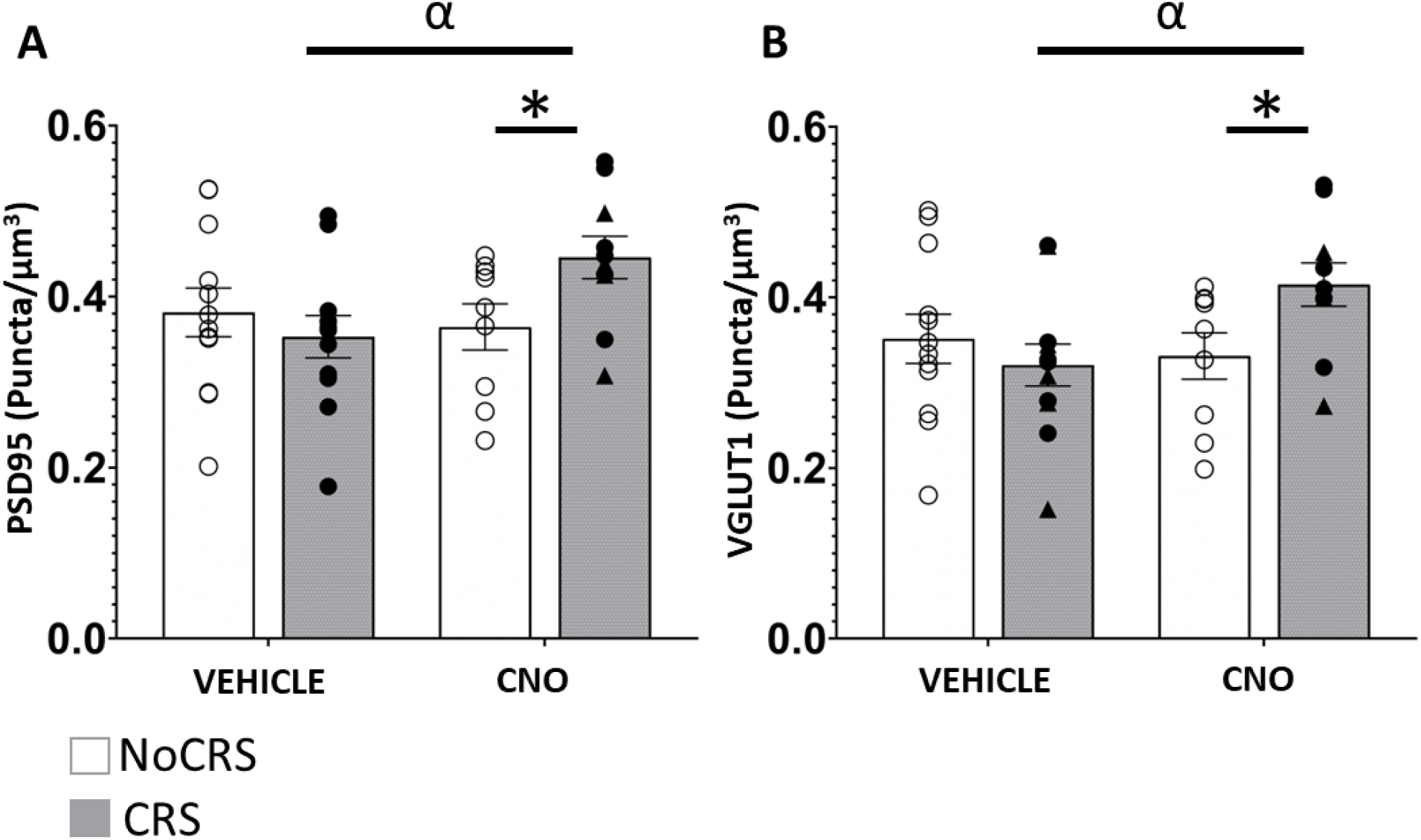
Effects of chronic hyperactivation of the amygdala combined or not with chronic stress on synaptic puncta density in the basolateral amygdala. Post-synaptic protein 95 (PSD95, **A**) and vesicular glutamate transporter 1 (VGLUT1, **B**) puncta density was quantified in the basolateral amygdala of mice treated with vehicle, clozapine-N-oxide (CNO) and/or chronic restraint stress (CRS). Value collected for each mouse as well as sex is represented (Δ for males and O for females). *p < 0.05, **p < 0.01 compared to vehicle treated group, α p<0.05 compared to CRS-exposed treated with vehicle group.

## 4. Discussion

The aim of this study was to determine whether cHOA is sufficient to induce depressive-like behavior and if it is a vulnerability factor to stress exposure. We found that mice with cHOA displayed a progressive increase in anxiety-like behavior observed at baseline in the phenotyper test. However, we showed that cHOA did not induce changes in anhedonia-like behavior measured using the sucrose consumption test every week. These results suggest that cHOA can induce anxiety-like but not depressive-like behavioral deficits. Animals were then subjected to CRS to determine if cHOA effects were exacerbated following stress exposure. We found similar behavioral deficits in mice exposed to CRS with or without cHOA in the phenotyper, NSF, and the sucrose consumption tests, suggesting that prior cHOA does not enhance response to chronic stress. Altogether, our results suggest that cHOA is sufficient to induce and sustain anxiety-like behavior, it is however not a susceptibility factor to stress. Morphological alterations associated with cHOA and CRS were then assessed using ex vivo MRI and synaptic puncta analysis. MRI results revealed that cHOA and CRS alone or combined did not alter the volume of key ROIs associated with emotion regulation; ROI volume did not correlate with behavioral emotionality. Lastly, analysis of synaptic puncta density revealed that CRS induces increases in PSD95 and VGLUT1 puncta density only in animals with prior cHOA; no correlational association between PSD95 and VGLUT1 puncta density changes, amygdala volume and behavioral emotionality were found. Altogether, MRI and synaptic puncta analysis suggest that cHOA alone or combined with 2 weeks CRS induce minor morphological changes unlike long term exposure to chronic stress.

Our study employed the DREADD approach with the purpose of selectively activating glutamatergic neurons in the BLA. Previous studies have used this approach to manipulate activity in various brain regions including the AMY (Aizenberg et al., 2019; Roth, 2016). Here we used micro-thin capillary needles to minimize the structural damage and allow for MRI analysis. Projection specific studies of the AMY have identified that acute activation of efferent glutamatergic projections from the BLA to the ventral hippocampus and medial PFC result in anxiogenic behavioral responses (Felix-Ortiz et al., 2013; Felix-Ortiz et al., 2016). Oppositely, stimulation of BLA projections to CeA and anterodorsal BNST induced anxiolytic behavior (Kim et al., 2013; Tye et al., 2011). However, these studies mostly used optogenetic approaches while chemogenetic acute activation of BLA glutamatergic neurons results was reported to cause an anxiogenic behavioral response (Siuda et al., 2016). These preclinical studies have mainly focused on understanding the role of amygdala activity and were limited to acute activation; the effects of chronic activation of the AMY were uninvestigated.

To our knowledge this is the first study to focus on examining the effects of chemogenetic cHOA on anxiety- and depressive-like behavior. We found that cHOA induced anxiety-like behavior at baseline in the phenotyper test confirming the effects observed acutely (Siuda et al., 2016). We also demonstrated chronic anxiety resulting from cHOA. This finding is relevant to the pathophysiology of anxiety disorder since various fMRI studies have considered that the hyperactivity of AMY is an indicator of clinical and subclinical anxiety (Etkin et al., 2004; Etkin and Wager, 2007). However, we also demonstrated that animals with cHOA did not subsequently develop anhedonia-like behavior. This suggests that cHOA alone is not sufficient to induced depressive-like behavioral deficits and is probably not an indicator of future symptom development. This is not supported by clinical observations suggesting that abnormal activation of the AMY measured using fMRI is a predictor of future onset of depressive symptoms (Swartz et al., 2015). However, it is in accordance with findings that amygdala hyperactivity can be a vital component of multiple anxiety associated disorders (PTSD, GAD, and OCD). Our results also suggest that despite the occurrence of anxiety and depressive disorder co-morbidities, chronic anxiety may rely independently from cHOA (or on additional mechanisms) to be able to lead to MDD. This is supported by evidence that healthy adults who have experienced early life stress with increased amygdala reactivity and do not become depressed may develop resiliency mechanisms (Yamamoto et al., 2017). Nevertheless, our results should be taken with caution since we only tested one behavior the sucrose consumption measuring anhedonia. Future studies should investigate broader behavioral assessments of anhedonia or helplessness to assert our conclusion.

Elevated amygdala activity seen in fMRI studies consistently occur following presentation of negative stimuli. This was shown in patients with anxiety disorders (ie. GAD, PTSD, OCD) and depression (Brühl et al., 2014b; Groenewold et al., 2013; Hayes et al., 2012; Swartz and Monk, 2014; Yang et al., 2010b). In addition, clinical research also showed that amygdala activation can be a vulnerability factor for future life stressors (Swartz et al., 2015). We thought to test this possibility using rodents. In our study we investigated if prior cHOA renders mice more susceptible to stress exposure. To test this hypothesis, we opted to combine cHOA with a chronic stress procedure that induced limited anhedonia- and anxiety-like behavior as well as minimal structural and morphological changes alone, Since we showed that CRS (2X 1hr/day protocol) for 2 weeks induced anxiety- and anhedonia like behavior and structural and morphological changes (Misquitta et al., 2021), we performed a less intensive CRS exposure protocol (1X/day). Using this approach, we found that mice with prior cHOA did not exhibit exacerbated response to CRS in anxiety- and depressive-like behaviors (as scored as behavioral emotionality) when behavior was observed after 3 or 15 days of CRS. The effects of CRS in mice with cHOA appeared somewhat greater in magnitude in the PT and overall z-emotionality score, suggesting that prior cHOA may slightly enhance stress effect on anxiety. Although not significant, this effect is in accordance with clinical findings of a link between increased amygdala reactivity in stressful situations in patients with anxiety disorders. For example, increase amygdala reactivity was associated with greater severity to stress response in soldiers who were exposed to stressful events during combat (Admon et al., 2009).

MRI volumetric changes associated with chronic stress have been studied across numerous rodent models (Anacker et al., 2016; Magalhães et al., 2018; Misquitta et al., 2021; Nikolova et al., 2018). A few studies have combined chemogenetic and fMRI approaches to investigate circuit and connectivity changes associated with acute activity manipulation of an ROI (Nakamura et al., 2020; Roelofs et al., 2017). Here, we aimed to examine potential association between behavioral, MRI and synaptic changes resulting from cHOA alone or combined with CRS. However, no significant volumetric changes were observed following these manipulations whether we considered specific limbic ROIs projecting to or target of the AMY, ROIs involved in MDD or the whole brain. We anticipated that CRS effects with the protocol used in this study (1X 1h/day) would result in little to no changes confirming results from a chronic stress study in rats (Magalhães et al., 2018). The lack of volumetric changes associated to seven weeks of cHOA suggests that hyperactivation of the AMY alone is not directly responsible for the volumetric changes associated with chronic stress. It is possible that dysfunction of multiple corticolimbic brain regions is required to recapitulate the effects of chronic stress. Interestingly, this result and the lack of correlation between AMY/AMY circuit ROIs and behavior also suggest that the chronic anxiety deficits we observed throughout the experiment did not rely on volumetric alterations of these brain regions. This may explain the mixed results between volume changes and anxiety symptoms expressed in anxiety disorders and MDD (Arnone et al., 2012a; Frodl et al., 2003; Hamilton et al., 2008). Finally, we found significant increases in both PSD95 and VGLUT1 puncta density in mice with cHOA subjected to chronic stress. These results are partially in line with literature showing that activation of neurons stimulates synaptogenesis (Andreae and Burrone, 2014). However, cHOA alone did not increase puncta density suggesting that other factors are involved in the hypertrophy of the AMY observed following stress (Misquitta et al., 2021; Vyas et al., 2004). In addition, we were unable to find any correlations between synaptic puncta density and volume or behavioral emotionality. This suggests that the elevated anxiety observed in cHOA animals did not rely on volumetric or synaptic changes and depend solely on the activity of the brain region. It is probably for this reason that acute activation of the AMY has the same effects (Siuda et al., 2016). On another note, studies have suggested that MRI-measuring volumetric changes rely on synaptic and dendritic loss or gain (Banasr et al., 2021). In the present study, we show that increases in puncta did not correlate with volume or was not associated with increased volume in cHOA animals subjected to chronic stress. This suggests that assumptions of the literature on MRI-volume, synapse or activity are not always true and need to be clearly demonstrated.

This study has several limitations. Firstly, we assess anxiety- and anhedonia-like behaviors using one test measuring each behavioral dimension. It is possible that the observed changes were specific to each test. Other tests measuring anxiety and anhedonia exist however most cannot be repeated hindering longitudinal analysis (Nollet et al., 2013; Planchez et al., 2019). Secondly, in our attempt to avoid ceiling effects of CRS and detect susceptibility, we used a less severe protocol for a shorter duration than our previous studies (Codeluppi et al., 2021; Misquitta et al., 2021; Prevot et al., 2019). It is possible that a longer CRS protocol is needed to observe greater susceptibility associated with cHOA. However we believe this is unlikely as the phenotyper test is very sensitive and is able to detect small variations in behavior (Prevot et al., 2019). Thirdly, we used two synaptic markers previously found to be associated with behavior and volume changes (Misquitta et al., 2021; Nikolova et al., 2018). An in-depth analysis of other synaptic architectural (ie. dendritic branching, synaptic button analysis) alterations might add to our understanding. Lastly, we had expected a much greater magnitude of change of MRI volumetric changes however the lack of any brain wide changes could be mainly due to activation of one small brain regions. Even though chronic stress exposure has detrimental effects to the amygdala it is not the only brain region impacted. Multiple chronic stress-based animal studies show evidence of MRI volumetric changes to affect several corticolimbic brain structures. Therefore, future studies should determine whether hypoactivity/hyperactivity in regions such as the mPFC and hippocampus, known to be involved in stress-related illnesses (Banasr et al., 2021), induces more extended volumetric changes.

In conclusion, this study investigated the behavioral, cellular and synaptic changes associated with chronic activation of the amygdala. We demonstrated its involvement in anxiety behavior and unlikely implicated in the development of depressive-like behavior and structural adaptations associated with chronic stress. This work provides information to clinical researchers that focuses on using AMY activity level or volume as a potential biomarker in stress-related illnesses such as GAD, PTSD and MDD.

## 7. Acknowledgements

We would also like to acknowledge the contributions of the CAMH animal facility personnel for animal care and genotyping services, specifically: Lori Dixon, Katrina Deverell, Kristen Fournier, and German Fernandes.

