## Supplement Material for "Behavioral and Neurostructural changes associated with Chronic Amygdala Hyperactivation"

Mounira Banasr^*^

**Table content**

**Supplementary Figure 1 (p2-3)**

**Supplementary Figure 2 (p4)**

**Supplementary Figure 3 (p5-6)**

**Supplementary Figure 4 (p7-8)**

**Supplementary Figure 5 (p9)**

**Supplementary Figure 6 (p10)**

**Supplementary Table 1	(p11)**

**Supplementary Table 2	(p12-16)**

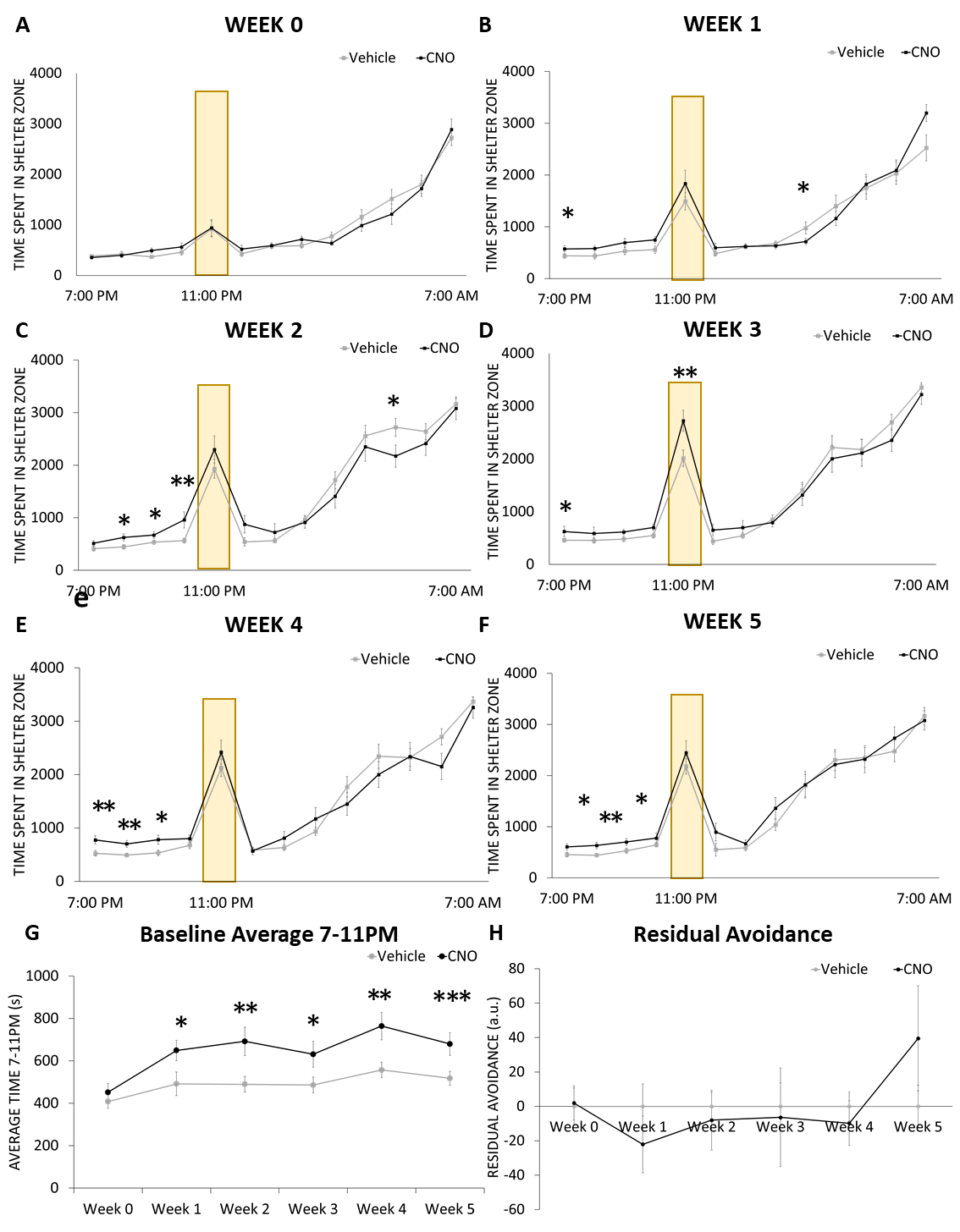

**Supplement Figure 1: Longitudinal effects of chronic hyperactivation of the amygdala on anxiety-like behavior in the phenotyper test (shelter zone).** Shelter zone time of mice treated or not with clozapine-N-oxide (CNO) was measured hourly between 7PM to 7AM in the PhenoTyper test. A white spotlight challenge occurs between 11-12PM (yellow bars). The time spent in the shelter zone prior to treatment (week 0, **A**) and every week of treatment (week 1, **B**; week 2 **C**, week 3, **D**; week 4, **E**; week 5, **F**) was assessed. Progressive changes in calculated baseline (7-11pm) averages and residual avoidance are shown in (**G)** and (**H)** respectively. Results are expressed as mean ± s.e.m. *p < 0.05, **p < 0.01, ***p<0.001 compared to vehicle treated group.

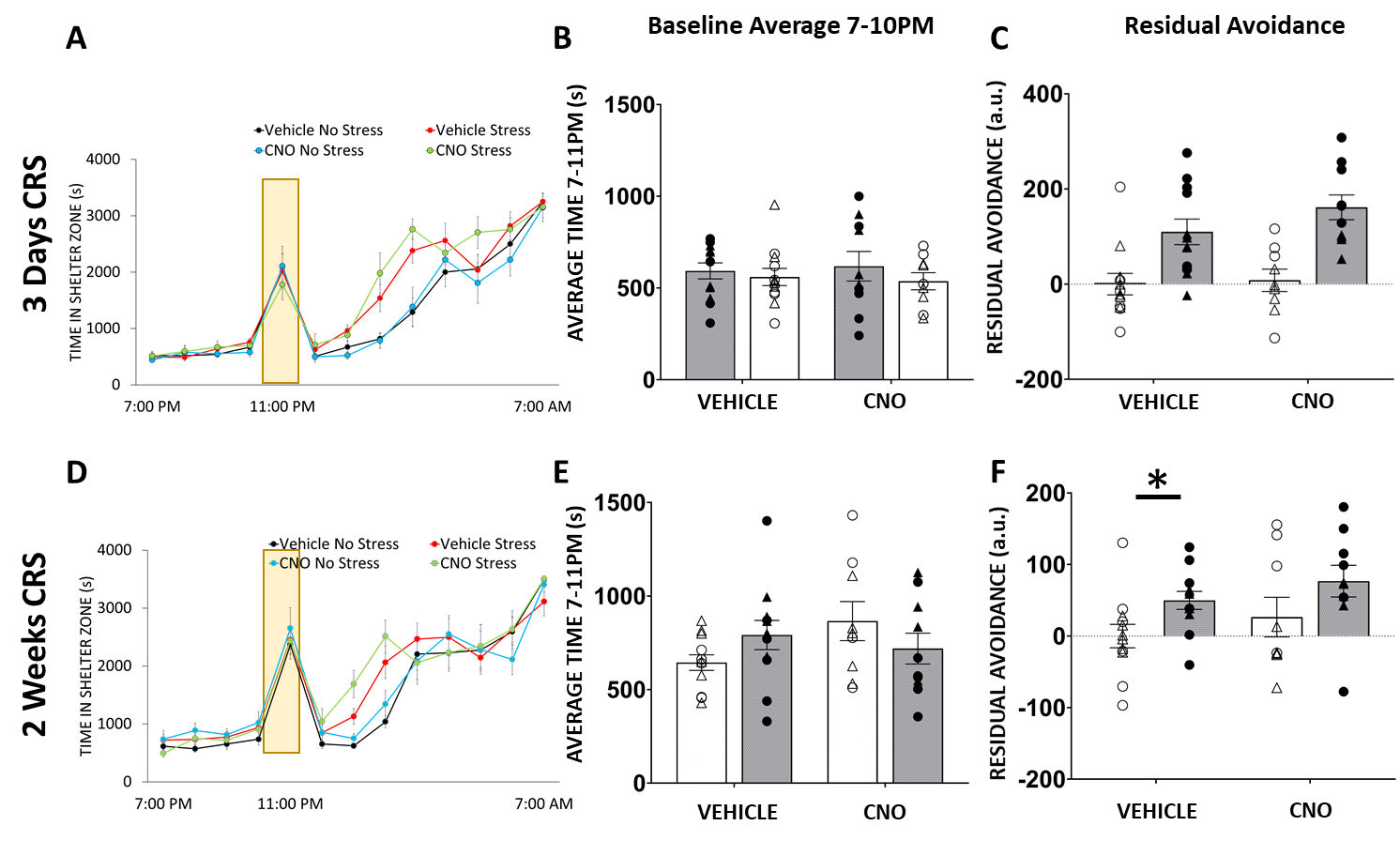

**Supplement Figure 2: Effects of chronic hyperactivation of the amygdala combined or not with chronic stress on anxiety-like behavior in the phenotyper test (shelter zone).** Shelter zone time of mice treated with vehicle, clozapine-N-oxide (CNO) and/or chronic restraint stress (CRS) was measured hourly between 7PM to 7AM in the PhenoTyper test on day 3 (**A**) and on day 15 of CRS exposure (**D**). A white spotlight challenge occurs between 11-12PM (yellow bars). Changes in calculated baseline (7-11pm) averages and residual avoidance found following 3 days of CRS are shown in (**B)** and (**C)** respectively. Baseline (7-11pm) averages (**E**) and residual avoidance (**F**) were also measured following 15 days of CRS. In B, C, E, F, Value collected for each mouse as well as sex is represented (△ for males and O for females). Results are expressed as mean ± s.e.m. *p < 0.05, compared to vehicle treated group.

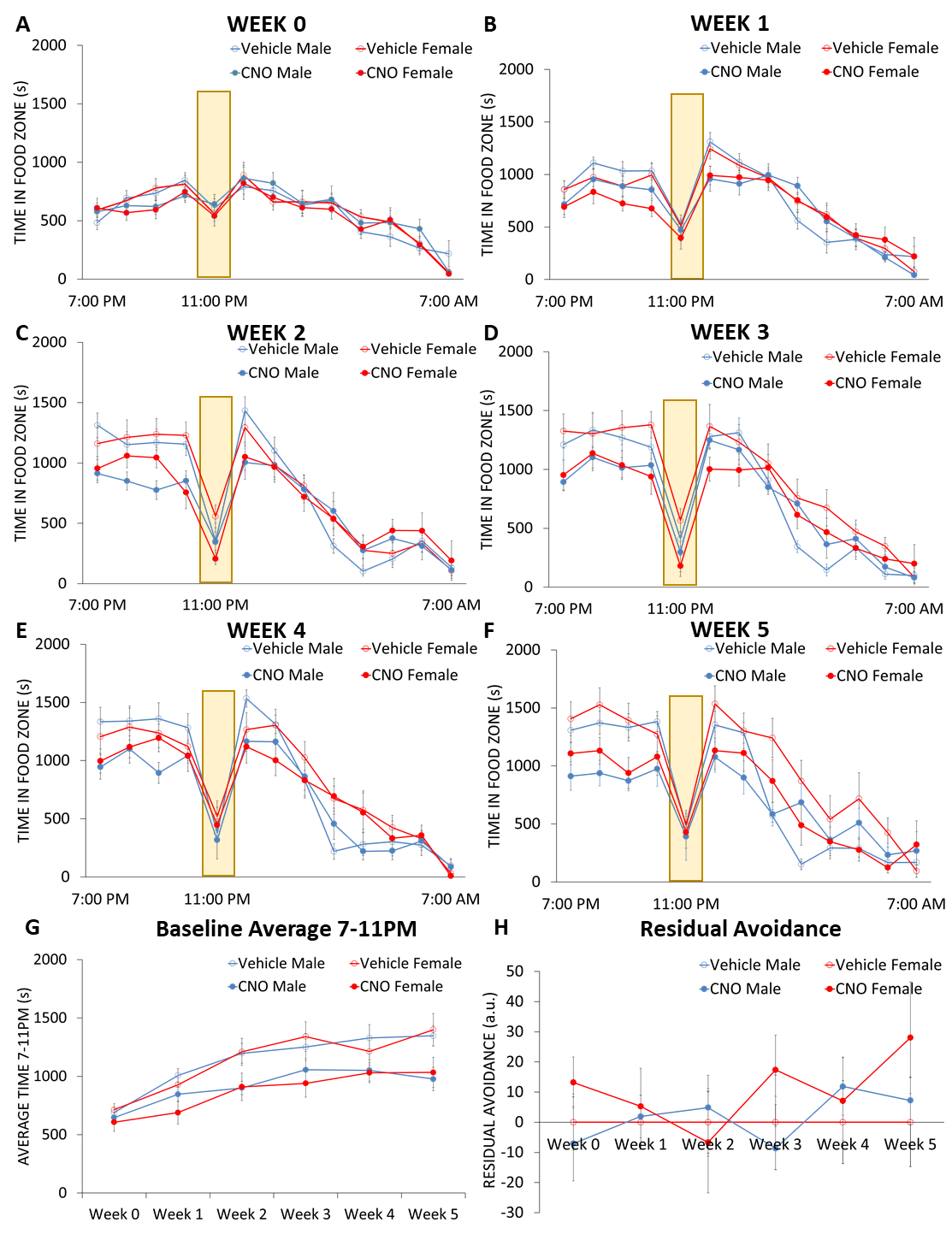

**Supplement Figure 3: Longitudinal effects of chronic hyperactivation of the amygdala on anxiety-like behavior in the phenotyper test (food zone) in males and females.** Food zone time of male and female mice treated or not with clozapine-N-oxide (CNO) was measured hourly between 7PM to 7AM in the PhenoTyper test. A white spotlight challenge occurs between 11-12PM (yellow bars). The time spent in the food zone prior to treatment (week 0, **A**) and every week of treatment (week 1, **B**; week 2 **C**, week 3, **D**; week 4, **E**; week 5, **F**) was assessed. Progressive changes in calculated baseline (7-11pm) averages and residual avoidance are shown in (**G)** and (**H)** respectively. Results are expressed as mean ± s.e.m.

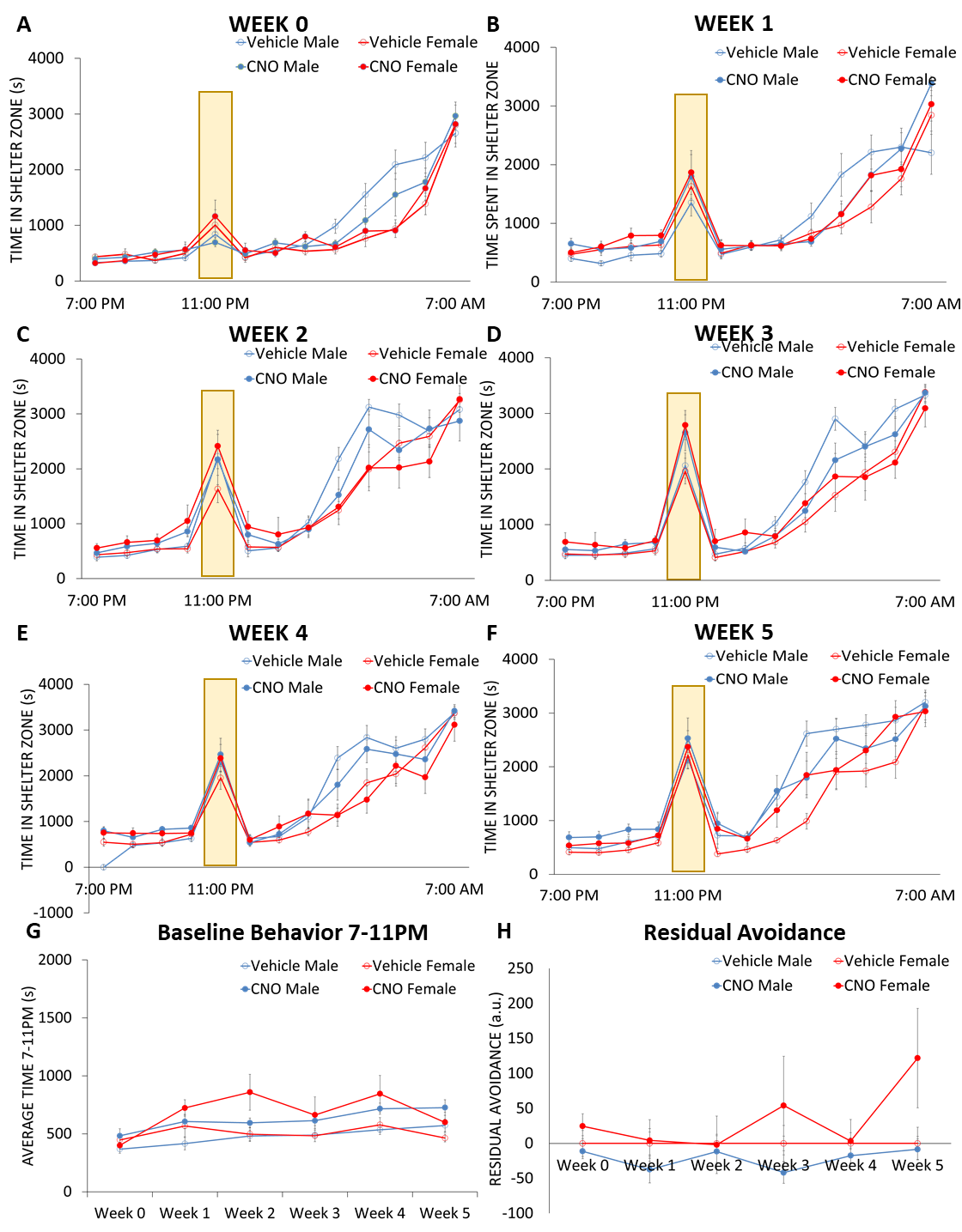

**Supplement Figure 4: Longitudinal effects of chronic hyperactivation of the amygdala on anxiety-like behavior in the phenotyper test (shelter zone) in males and females.** Shelter zone time of male and female mice treated or not with clozapine-N-oxide (CNO) was measured hourly between 7PM to 7AM in the PhenoTyper test. A white spotlight challenge occurs between 11-12PM (yellow bars). The time spent in the shelter zone prior to treatment (week 0, **A**) and every week of treatment (week 1, **B**; week 2 **C**, week 3, **D**; week 4, **E**; week 5, **F**) was assessed. Progressive changes in calculated baseline (7-11pm) averages and residual avoidance are shown in (**G)** and (**H)** respectively. Results are expressed as mean ± s.e.m.

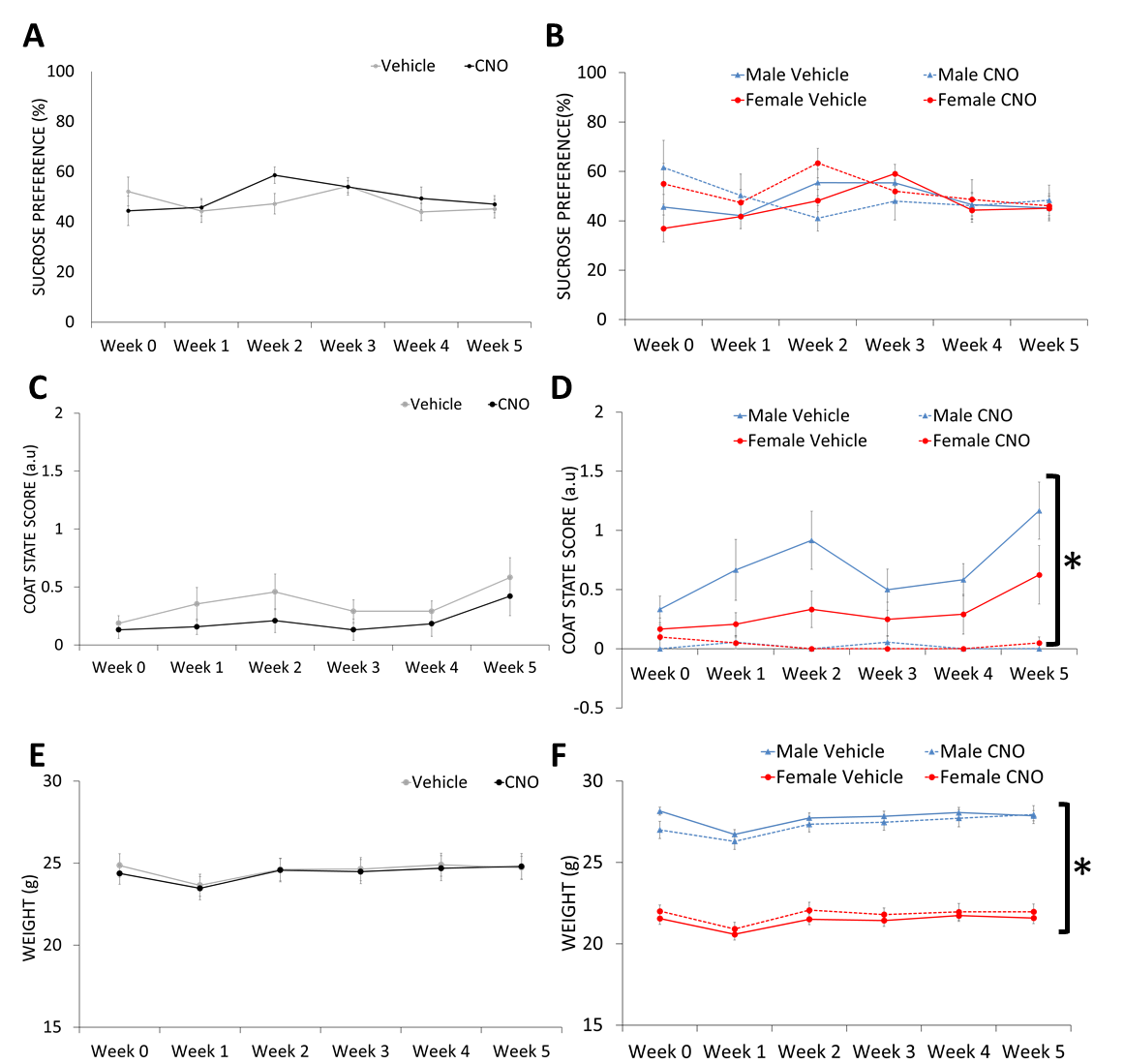

**Supplement Figure 5: Longitudinal effects of chronic hyperactivation of the amygdala on sucrose preference, coat state and weight.** Mice treated with vehicle or clozapine-N-oxide (CNO) were tested weekly the sucrose preference test and coat state assessment. Mice were also weighted every week. Graphs show data collected before treatment week 0 and every following week for 5 weeks for all animals split by group (**A**: sucrose preference; **C**: coat state; **E**: weight) or split by group and sex (**B**: sucrose preference; **D**: coat state; **F**: weight) Overall z-emotionality scores calculated with respect to average control group are shown in (**D**). Results are expressed as mean ± s.e.m. *p < 0.05, main effect of sex.

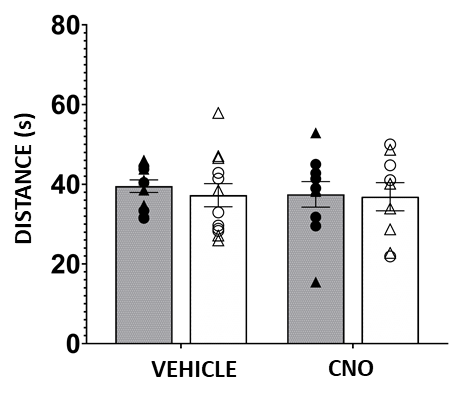

**Supplementary Figure 6: Effects of chronic hyperactivation of the amygdala combined or not with chronic stress on locomotor activity.** Distance travelled (in meters) by mice treated with vehicle or clozapine-N-oxide (CNO) and/or subjected chronic restraint stress (CRS) was measured for 30min in a home-cage like arena. to were tested weekly the sucrose preference test and coat state assessment. Value collected for each mouse as well as sex is represented (△ for males and O for females). Results are expressed as mean ± s.e.m.

Table 1: Analysis of variance of 26 MDD-associated brain region volumes and Pearson correlation with z-emotionality score (adjusted for multiple comparisons).

| **MDD Associated ROI** | **Group (p-value)** | **Stress (p-value)** |  | **Pearson R** | **p Value** | **q Value** |
| --- | --- | --- | --- | --- | --- | --- |
| Amygdala | 0.87 | 0.34 |  | 0.24 | 0.13 | 0.89 |
| Globus Pallidus | 0.72 | 0.43 |  | 0.14 | 0.36 | 0.89 |
| Hypothalamus | 0.56 | 0.17 |  | 0.17 | 0.28 | 0.89 |
| Midbrain | 0.36 | 0.54 |  | 0.20 | 0.20 | 0.89 |
| Nucleus Accumbens | 0.54 | 0.90 |  | 0.12 | 0.45 | 0.89 |
| Striatum | 0.47 | 0.26 |  | 0.11 | 0.50 | 0.89 |
| Thalamus | 0.89 | 0.08 |  | 0.13 | 0.39 | 0.89 |
| Cingulate Cortex Area 24A | 0.18 | 0.13 |  | 0.16 | 0.31 | 0.89 |
| Cingulate Cortex Area 24B | 0.28 | 0.31 |  | -0.10 | 0.54 | 0.89 |
| Cingulate Cortex Area 25 | 0.33 | 0.03 |  | 0.28 | 0.07 | 0.89 |
| Cingulate Cortex Area 29a | 0.85 | 0.42 |  | 0.03 | 0.85 | 0.89 |
| Cingulate Cortex Area 29b | 0.34 | 0.30 |  | 0.03 | 0.84 | 0.89 |
| Cingulate Cortex Area 29c | 0.24 | 0.99 |  | -0.15 | 0.34 | 0.89 |
| Cingulate Cortex Area 30 | 0.34 | 0.82 |  | -0.03 | 0.85 | 0.89 |
| Cingulate Cortex Area 32 | 0.98 | 0.50 |  | 0.05 | 0.77 | 0.89 |
| Dorsolateral Orbital Cortex | 0.23 | 0.12 |  | -0.09 | 0.56 | 0.89 |
| Frontal Association Cortex | 0.23 | 0.40 |  | -0.10 | 0.51 | 0.89 |
| Insular Region Not Subdivided | 0.85 | 0.35 |  | -0.04 | 0.82 | 0.89 |
| Lateral Orbital Cortex | 0.26 | 0.24 |  | -0.09 | 0.55 | 0.89 |
| Lateral Parietal Association Cortex | 0.53 | 0.42 |  | 0.08 | 0.63 | 0.89 |
| Medial Orbital Cortex | 0.40 | 0.20 |  | 0.01 | 0.94 | 0.94 |
| Medial Parietal Association Cortex | 0.35 | 0.55 |  | 0.04 | 0.80 | 0.89 |
| Temporal Aassociation Area | 0.54 | 0.70 |  | -0.04 | 0.80 | 0.89 |
| Ventral Orbital Cortex | 0.96 | 0.60 |  | 0.04 | 0.78 | 0.89 |
| Dentate Gyrus of the Hippocampus | 0.61 | 0.51 |  | 0.12 | 0.45 | 0.89 |
| Hippocampal Formation | 0.78 | 0.92 |  | 0.08 | 0.62 | 0.89 |

Table 2: Brain wide volumetric analysis of mice exposed to cHOA and CRS exposure as compared to vehicle mice with no stress.

| **Regions of Interest** | **Group (p-value)** | **q value** | **Stress (p-value)** | **q value** |
| --- | --- | --- | --- | --- |
| Frontal_cortex_area_3 | 0.00 | 0.62 | 0.78 | 0.94 |
| cuneate_nucleus | 0.01 | 0.62 | 0.45 | 0.92 |
| Primary_somatosensory_cortex | 0.01 | 0.62 | 0.79 | 0.94 |
| Medial_preoptic_nucleus | 0.03 | 0.99 | 0.08 | 0.92 |
| Primary_somatosensory_cortex_upper_lip_region | 0.03 | 0.99 | 0.50 | 0.94 |
| nucleus_interpositus | 0.03 | 0.99 | 0.12 | 0.92 |
| cerebral_aqueduct | 0.06 | 0.99 | 0.33 | 0.92 |
| pons | 0.06 | 0.99 | 0.72 | 0.94 |
| Cingulate_cortex_area_24a.1 | 0.06 | 0.99 | 0.08 | 0.92 |
| anterior_commissure_pars_anterior | 0.08 | 0.99 | 0.13 | 0.92 |
| colliculus_superior | 0.08 | 0.99 | 0.92 | 0.98 |
| lateral_olfactory_tract | 0.09 | 0.99 | 0.07 | 0.92 |
| Primary_somatosensory_cortex_jaw_region | 0.09 | 0.99 | 0.62 | 0.94 |
| facial_nerve_cranial_nerve_7 | 0.10 | 0.99 | 0.55 | 0.94 |
| cerebellar_peduncle_inferior | 0.11 | 0.99 | 0.78 | 0.94 |
| subependymale_zone_rhinocele | 0.11 | 0.99 | 0.08 | 0.92 |
| olfactory_peduncle | 0.12 | 0.99 | 0.34 | 0.92 |
| paramedian_lobule | 0.12 | 0.99 | 0.10 | 0.92 |
| crus_2_white_matter | 0.13 | 0.99 | 0.67 | 0.94 |
| Cingulum | 0.13 | 0.99 | 0.09 | 0.92 |
| pontine_nucleus | 0.14 | 0.99 | 0.17 | 0.92 |
| fourth_ventricle | 0.16 | 0.99 | 0.90 | 0.98 |
| crus_1_ansiform_lobule_lobule_6 | 0.16 | 0.99 | 0.24 | 0.92 |
| mammilothalamic_tract | 0.17 | 0.99 | 0.10 | 0.92 |
| medial_septum | 0.18 | 0.99 | 0.83 | 0.94 |
| lobule_3_central_lobule_dorsal | 0.19 | 0.99 | 0.81 | 0.94 |
| lobule_10_nodulus | 0.21 | 0.99 | 0.29 | 0.92 |
| lateral_ventricle | 0.21 | 0.99 | 0.63 | 0.94 |
| Secondary_visual_cortex_mediolateral_area | 0.22 | 0.99 | 0.54 | 0.94 |
| Accessory_olfactory_bulb_glomerular_external_plexiform_and_mitral_cell_layer | 0.22 | 0.99 | 0.93 | 0.98 |
| Secondary_visual_cortex_mediomedial_area | 0.22 | 0.99 | 0.75 | 0.94 |
| periaqueductal_grey | 0.22 | 0.99 | 0.32 | 0.92 |
| Frontal_association_cortex | 0.23 | 0.99 | 0.36 | 0.92 |
| Dorsolateral_orbital_cortex | 0.23 | 0.99 | 0.14 | 0.92 |
| fasciculus_retroflexus | 0.23 | 0.99 | 0.01 | 0.92 |
| optic_tract | 0.24 | 0.99 | 0.93 | 0.98 |
| dentate_nucleus | 0.24 | 0.99 | 0.34 | 0.92 |
| Primary_somatosensory_cortex_barrel_field | 0.24 | 0.99 | 0.65 | 0.94 |
| medulla | 0.24 | 0.99 | 0.78 | 0.94 |
| Cingulate_cortex_area_29c | 0.24 | 0.99 | 0.99 | 0.99 |
| Cingulate_cortex_area_24b | 0.25 | 0.99 | 0.41 | 0.92 |
| CA3Py_Inner | 0.26 | 0.99 | 0.69 | 0.94 |
| lobule_10_white_matter | 0.27 | 0.99 | 0.38 | 0.92 |
| Lateral_orbital_cortex | 0.27 | 0.99 | 0.23 | 0.92 |
| anterior_lobule_white_matter | 0.28 | 0.99 | 0.76 | 0.94 |
| crus_2_ansiform_lobule_lobule_7 | 0.30 | 0.99 | 0.65 | 0.94 |
| paraflocculus_white_matter | 0.30 | 0.99 | 0.18 | 0.92 |
| anterior_commissure_pars_posterior | 0.31 | 0.99 | 0.50 | 0.94 |
| fimbria | 0.32 | 0.99 | 0.72 | 0.94 |
| Olfactory_bulb_glomerular_layer | 0.32 | 0.99 | 0.44 | 0.92 |
| trunk_of_arbor_vita | 0.32 | 0.99 | 0.21 | 0.92 |
| fastigial_nucleus | 0.33 | 0.99 | 0.09 | 0.92 |
| Primary_motor_cortex | 0.33 | 0.99 | 0.79 | 0.94 |
| Cingulate_cortex_area_25 | 0.34 | 0.99 | 0.04 | 0.92 |
| Secondary_somatosensory_cortex | 0.34 | 0.99 | 0.80 | 0.94 |
| Cingulate_cortex_area_29b | 0.34 | 0.99 | 0.27 | 0.92 |
| Claustrum | 0.34 | 0.99 | 0.38 | 0.92 |
| Cingulate_cortex_area_24a | 0.34 | 0.99 | 0.27 | 0.92 |
| Cingulate_cortex_area_30 | 0.34 | 0.99 | 0.74 | 0.94 |
| midbrain | 0.35 | 0.99 | 0.64 | 0.94 |
| Medial_parietal_association_cortex | 0.36 | 0.99 | 0.57 | 0.94 |
| CA10r | 0.37 | 0.99 | 0.73 | 0.94 |
| Anterior_olfactory_nucleus | 0.38 | 0.99 | 1.00 | 1.00 |
| Primary_visual_cortex_monocular_area | 0.39 | 0.99 | 0.51 | 0.94 |
| Medial_orbital_cortex | 0.40 | 0.99 | 0.17 | 0.92 |
| colliculus_inferior | 0.40 | 0.99 | 0.11 | 0.92 |
| trunk_of_simple_and_crus_1_white_matter | 0.41 | 0.99 | 0.78 | 0.94 |
| anterior_lobule_lobules_4_5 | 0.42 | 0.99 | 0.71 | 0.94 |
| Olfactory_bulb_granule_cell_layer | 0.43 | 0.99 | 0.47 | 0.92 |
| Accessory_olfactory_bulb_granule_cell_layer | 0.43 | 0.99 | 0.47 | 0.92 |
| Primary_somatosensory_cortex_forelimb_region | 0.43 | 0.99 | 0.97 | 0.99 |
| trunk_of_crus_2_and_paramedian_white_matter | 0.44 | 0.99 | 0.19 | 0.92 |
| corpus_callosum | 0.44 | 0.99 | 0.81 | 0.94 |
| Secondary_motor_cortex | 0.44 | 0.99 | 0.64 | 0.94 |
| ventral_tegmental_decussation | 0.44 | 0.99 | 0.29 | 0.92 |
| MoDG | 0.44 | 0.99 | 0.69 | 0.94 |
| striatum | 0.46 | 0.99 | 0.20 | 0.92 |
| cerebellar_peduncle_middle | 0.46 | 0.99 | 0.30 | 0.92 |
| habenular_commissure | 0.47 | 0.99 | 0.20 | 0.92 |
| Cingulate_cortex_area_24b.1 | 0.47 | 0.99 | 0.33 | 0.92 |
| Temporal_association_area | 0.48 | 0.99 | 0.47 | 0.92 |
| fundus_of_striatum | 0.48 | 0.99 | 0.45 | 0.92 |
| Claustrum_dorsal_part | 0.49 | 0.99 | 0.42 | 0.92 |
| Intermediate_nucleus_of_the_endopiriform_claustrum | 0.50 | 0.99 | 0.81 | 0.94 |
| lateral_septum | 0.50 | 0.99 | 0.50 | 0.94 |
| Secondary_auditory_cortex_ventral_area | 0.50 | 0.99 | 0.44 | 0.92 |
| Primary_somatosensory_cortex_shoulder_region | 0.51 | 0.99 | 0.75 | 0.94 |
| hypothalamus | 0.51 | 0.99 | 0.21 | 0.92 |
| copula_white_matter | 0.52 | 0.99 | 0.15 | 0.92 |
| stria_medullaris | 0.53 | 0.99 | 0.12 | 0.92 |
| Lateral_parietal_association_cortex | 0.53 | 0.99 | 0.46 | 0.92 |
| nucleus_accumbens | 0.54 | 0.99 | 0.82 | 0.94 |
| crus_1_white_matter | 0.55 | 0.99 | 0.18 | 0.92 |
| cerebellar_peduncle_superior | 0.55 | 0.99 | 0.28 | 0.92 |
| Caudomedial_entorhinal_cortex | 0.56 | 0.99 | 0.67 | 0.94 |
| superior_olivary_complex | 0.58 | 0.99 | 0.56 | 0.94 |
| lobule_9_white_matter | 0.59 | 0.99 | 0.19 | 0.92 |
| simple_lobule_lobule_6 | 0.59 | 0.99 | 0.58 | 0.94 |
| third_ventricle | 0.59 | 0.99 | 0.98 | 0.99 |
| DG | 0.59 | 0.99 | 0.62 | 0.94 |
| Dorsolateral_entorhinal_cortex | 0.60 | 0.99 | 0.18 | 0.92 |
| subiculum | 0.60 | 0.99 | 0.95 | 0.99 |
| lobule_8_white_matter | 0.60 | 0.99 | 0.35 | 0.92 |
| lobule_9_uvula | 0.61 | 0.99 | 0.33 | 0.92 |
| mammillary_bodies | 0.61 | 0.99 | 0.56 | 0.94 |
| Primary_somatosensory_cortex_dysgranular_zone | 0.63 | 0.99 | 0.73 | 0.94 |
| CA3Py_Outer | 0.63 | 0.99 | 0.30 | 0.92 |
| Ectorhinal_cortex | 0.64 | 0.99 | 0.63 | 0.94 |
| Dorsal_tenia_tecta | 0.64 | 0.99 | 0.22 | 0.92 |
| paraflocculus_PFL | 0.66 | 0.99 | 0.28 | 0.92 |
| Claustrum_ventral_part | 0.66 | 0.99 | 0.91 | 0.98 |
| trunk_of_lobules_6_8_white_matter | 0.67 | 0.99 | 0.11 | 0.92 |
| bed_nucleus_of_stria_terminalis | 0.69 | 0.99 | 0.17 | 0.92 |
| copula_pyramis_lobule_8 | 0.69 | 0.99 | 0.31 | 0.92 |
| Dorsal_intermediate_entorhinal_cortex | 0.70 | 0.99 | 0.31 | 0.92 |
| Olfactory_bulb_internal_plexiform_layer | 0.71 | 0.99 | 0.17 | 0.92 |
| CA1 | 0.72 | 0.99 | 0.96 | 0.99 |
| paramedian_lobule_lobule_7 | 0.72 | 0.99 | 0.24 | 0.92 |
| globus_pallidus | 0.72 | 0.99 | 0.44 | 0.92 |
| lobule_6_declive | 0.73 | 0.99 | 0.79 | 0.94 |
| lobules_6_7_white_matter | 0.73 | 0.99 | 0.72 | 0.94 |
| Ventral_tenia_tecta | 0.74 | 0.99 | 0.84 | 0.94 |
| GrDG | 0.74 | 0.99 | 0.60 | 0.94 |
| lobules_4_5_culmen_ventral_and_dorsal | 0.74 | 0.99 | 0.83 | 0.94 |
| pre_para_subiculum | 0.74 | 0.99 | 0.71 | 0.94 |
| internal_capsule | 0.75 | 0.99 | 0.73 | 0.94 |
| flocculus_white_matter | 0.75 | 0.99 | 0.72 | 0.94 |
| lobule_3_white_matter | 0.75 | 0.99 | 0.59 | 0.94 |
| stria_terminalis | 0.75 | 0.99 | 0.73 | 0.94 |
| Primary_visual_cortex_binocular_area | 0.75 | 0.99 | 0.66 | 0.94 |
| Olfactory_bulb_external_plexiform_layer | 0.76 | 0.99 | 0.21 | 0.92 |
| lobules_1_2_lingula_and_central_lobule_ventral | 0.76 | 0.99 | 0.79 | 0.94 |
| lobules_4_5_white_matter | 0.76 | 0.99 | 0.92 | 0.98 |
| SLu | 0.76 | 0.99 | 0.30 | 0.92 |
| LMol | 0.78 | 0.99 | 0.81 | 0.94 |
| trunk_of_lobules_1_3_white_matter | 0.79 | 0.99 | 0.31 | 0.92 |
| CA1Py | 0.80 | 0.99 | 0.89 | 0.98 |
| Piriform_cortex | 0.80 | 0.99 | 0.42 | 0.92 |
| olfactory_tubercle | 0.82 | 0.99 | 0.91 | 0.98 |
| interpedunclar_nucleus | 0.82 | 0.99 | 0.06 | 0.92 |
| medial_lemniscus_medial_longitudinal_fasciculus | 0.82 | 0.99 | 0.38 | 0.92 |
| Secondary_visual_cortex_lateral_area | 0.83 | 0.99 | 0.96 | 0.99 |
| Primary_auditory_cortex | 0.84 | 0.99 | 0.53 | 0.94 |
| posterior_commissure | 0.85 | 0.99 | 0.10 | 0.92 |
| PoDG | 0.85 | 0.99 | 0.45 | 0.92 |
| Cingulate_cortex_area_29a | 0.85 | 0.99 | 0.40 | 0.92 |
| CA3 | 0.85 | 0.99 | 0.32 | 0.92 |
| Rostral_amygdalopiriform_area | 0.85 | 0.99 | 0.98 | 0.99 |
| Insular_region_not_subdivided | 0.85 | 0.99 | 0.34 | 0.92 |
| Medial_entorhinal_cortex | 0.85 | 0.99 | 0.38 | 0.92 |
| Amygdalopiriform_transition_area | 0.86 | 0.99 | 0.41 | 0.92 |
| amygdala | 0.86 | 0.99 | 0.27 | 0.92 |
| CA30r | 0.86 | 0.99 | 0.30 | 0.92 |
| thalamus | 0.88 | 0.99 | 0.05 | 0.92 |
| Posterolateral_cortical_amygdaloid_area | 0.88 | 0.99 | 0.72 | 0.94 |
| Medial_amygdala | 0.88 | 0.99 | 0.29 | 0.92 |
| corticospinal_tract_pyramids | 0.89 | 0.99 | 0.97 | 0.99 |
| lobule_8_pyramis | 0.90 | 0.99 | 0.64 | 0.94 |
| Cortex_amygdala_transition_zones | 0.90 | 0.99 | 0.54 | 0.94 |
| CA20r | 0.90 | 0.99 | 0.25 | 0.92 |
| Posteromedial_cortical_amygdaloid_area | 0.91 | 0.99 | 0.18 | 0.92 |
| Dorsal_nucleus_of_the_endopiriform | 0.91 | 0.99 | 0.83 | 0.94 |
| Parietal_cortex_posterior_area_rostral_part | 0.92 | 0.99 | 0.74 | 0.94 |
| Primary_somatosensory_cortex_hindlimb_region | 0.92 | 0.99 | 0.64 | 0.94 |
| CA3Rad | 0.93 | 0.99 | 0.41 | 0.92 |
| CA2Py | 0.93 | 0.99 | 0.31 | 0.92 |
| Primary_somatosensory_cortex_trunk_region | 0.93 | 0.99 | 0.72 | 0.94 |
| fornix | 0.93 | 0.99 | 0.22 | 0.92 |
| Ventral_intermediate_entorhinal_cortex | 0.93 | 0.99 | 0.45 | 0.92 |
| CA2Rad | 0.93 | 0.99 | 0.56 | 0.94 |
| flocculus_FL | 0.93 | 0.99 | 0.47 | 0.92 |
| Primary_visual_cortex | 0.93 | 0.99 | 0.53 | 0.94 |
| Perirhinal_cortex | 0.95 | 0.99 | 0.91 | 0.98 |
| basal_forebrain | 0.95 | 0.99 | 0.77 | 0.94 |
| Secondary_auditory_cortex_dorsal_area | 0.95 | 0.99 | 0.89 | 0.98 |
| Ventral_orbital_cortex | 0.96 | 0.99 | 0.59 | 0.94 |
| cerebral_peduncle | 0.97 | 0.99 | 0.94 | 0.99 |
| simple_lobule_white_matter | 0.97 | 0.99 | 0.33 | 0.92 |
| lobule_1_2_white_matter | 0.97 | 0.99 | 0.44 | 0.92 |
| Cingulate_cortex_area_32 | 0.97 | 0.99 | 0.42 | 0.92 |
| CA1Rad | 0.97 | 0.99 | 0.80 | 0.94 |
| Olfactory_bulb_mitral_cell_layer | 0.98 | 0.99 | 0.21 | 0.92 |
| lobule_7_tuber_or_folium | 0.98 | 0.99 | 0.73 | 0.94 |
| inferior_olivary_complex | 0.98 | 0.99 | 0.43 | 0.92 |
| CA2 | 0.98 | 0.99 | 0.37 | 0.92 |
| Ventral_nucleus_of_the_endopiriform_claustrum | 1.00 | 1.00 | 0.60 | 0.94 |
